# NSUN2 Facilitates DICER Cleavage of DNA Damage-Associated R-Loops to Promote Repair

**DOI:** 10.1101/2024.04.30.591877

**Authors:** Adele Alagia, Arianna Di Fazio, Kamal Ajit, Qilin Long, Monika Gullerova

## Abstract

DNA integrity is constantly challenged by both endogenous and exogenous damaging agents, resulting in various forms of damage. Failure to repair DNA accurately leads to genomic instability, a hallmark of cancer. Distinct pathways exist to repair different types of DNA damage. Double-strand breaks (DSBs) represent particularly severe form of damage, due to the physical separation of DNA strands. The repair of DSBs requires the activity of RNA Polymerase II (RNAPII) and the generation of Damage-associated transcripts (DARTs).

Here we show that the RNA m5C-methyltransferase NSUN2 localizes to DSBs in a transcription-dependent manner, where it binds to and methylates DARTs. The depletion of NSUN2 results in an accumulation of nascent primary DARTs around DSBs. Furthermore, we detected an RNA-dependent interaction between NSUN2 and DICER, which was stimulated by DNA damage. NSUN2 activity promoted DICER cleavage of DARTs-associated R-loops, which is required for efficient DNA repair.

We report a previously unrecognized role of the RNA m5C-methyltransferase NSUN2 within the RNA-dependent DNA damage response, highlighting its function as a DICER chaperone for the clearance of non-canonical substrates such as DARTs, thereby contributing to genomic integrity.

## Main

Genetic information, encoded in DNA, is constantly exposed to environmental or endogenous damaging agents, leading to numerous damaged sites within the genome. To preserve DNA integrity, cells have developed proficient damage response and repair mechanisms^1^. Failure or errors in DNA repair can result in the accumulation of damage, which negatively affects cellular viability and contributes to diseases, particularly cancer^2^. Double-Strand Breaks (DSBs), caused by the simultaneous breakage of the DNA double helix’s both strands, represent the most toxic form of DNA damage. DSBs are primarily repaired through the rapid and error-prone process of non-homologous end joining (NHEJ) or the more accurate homologous recombination (HR)^3^.

The activation of the DNA Damage Response (DDR) involves both protein and RNA molecules. Recent evidence points to the role of various long non-coding RNAs in RNA-dependent DNA repair (RDDR), acting in *cis* or *trans* ^4^.

DSBs can serve as functional promoters, triggering *de novo* transcription that generates long non-coding transcripts crucial for efficient DNA repair. In particular, the MRN complex coordinates the preinitiation complex (PIC) formation at DSB ends, which facilitates the recruitment of active RNA Polymerase II (RNAPII) to initiate bidirectional transcription^5, 6^. This process results in the generation of damage-induced long non coding RNAs (dilncRNAs), which help to recruit various DDR factors^7^. Moreover, dilncRNAs can directly facilitate DDR by promoting end resection, a critical step in HR repair. We showed previously that c-Abl mediated phosphorylation of the Y1 CTD RNAPII, induced *de novo* transcription at DSBs. Upon DNA damage, Y1P RNAPII transcribes away from DSBs, producing long transcripts (primary DART) that hybridize with their template DNA to form R-loops. This action initiates antisense transcription directed toward the DSB (secondary DART), leading to the formation of double-stranded RNAs (dsRNAs). DARTs are crucial in the recruitment of DDR factors, such as 53BP1 and MDC1, thereby facilitating effective DDR^8^.

Interestingly, dilncRNs and DARTs can be processed into small non-coding RNAs, known as DNA damage response RNAs (DDRNAs), which have direct roles in DDR. The depletion of DICER and DROSHA, key RNA processing enzymes in RNA interference (RNAi), significantly reduces the formation of foci containing DDR factors such as 53BP1, pATM, and MDC1^9–12^. Notably, the knockdown of other factors involved in miRNA biogenesis, including the three GW182-like proteins, does not affect DDR foci formation, indicating a miRNA-independent role for DROSHA and DICER in DDR.

Post-transcriptional RNA modifications, including N6-methyladenosine (m6A), N1-methyladenosine (m1A), 5-methylcytosine (m5C), 2′-O-methylation (2′-OMe), and pseudouridine (Ψ), play crucial roles in regulating various aspects of RNA biology, including stability, splicing, localization, and translation. These modifications are dynamically regulated by “writers,” “readers,” and “erasers” enzymes. Each of these enzymes contributes to the modification process, ensuring proper cellular function. Furthermore, RNA modifications have been linked to DDR pathways, highlighting their significance in the maintenance of genome integrity ^13, 14^. For example, m6A RNA methylation facilitates the recruitment of Pol κ to UV-induced DNA lesions, thus accelerating repair^15^. Moreover, m6A RNA modifications, mediated by METTL3 and recognized by YTHDC1, accumulate at DSBs to promote HR^16^. Likewise, m6A modifications on RNA within DNA:RNA hybrids at DSBs modulates their stability^16^. Similarly, m5C accumulates at damaged sites and contributes to DNA repair, particularly within the context of R-loops. The recruitment of the writer enzyme TRDMT1 to DNA damage sites is crucial for m5C-mediated repair, as its deficiency hinders HR ^17^. NSUN2, another 5C methyltransferase, is primarily known for tRNA-methylation, but has also been implicated in modifying other RNA substrates, such as mRNA and long non coding RNA (lncRNA)^18–21^. NSUN2’s amplification/overexpression is common in many cancers^22^. Therefore, the understanding of role NSUN2’s role in DSBs repair is of great importance.

Here we show that NSUN2 is localized at DSBs in transcription-dependent manner. Moreover, using a modified FISH-PLA technique ^23^, we show that NSUN2 binds to and methylates DARTs. NSUN2 depletion results in an accumulation of nascent RNA around DSBs, particularly primary DARTs. Furthermore, we detected an RNA-dependent interaction between NSUN2 and DICER, stimulated by DNA damage. NSUN2 activity facilitates DICER-mediated cleavage of DARTs-associated R-loops, essential for efficient DNA repair.

We propose a novel role for the m5C-methyltransferase NSUN2 within RDDR. Our findings provide insights into the function of NSUN2 as a DICER co-factor for the cleavage and clearance of non-canonical substrates such as DARTs.

## Results

### NSUN2 is recruited to double strand breaks

A previous study on scaRNA2 and DNA damage repair revealed its role in stimulating HR-mediated DNA damage response^24^. Additionally, a miCLIP assay identified scaRNA2 as a substrate for NSUN2-mediated m5C modification^25^. To investigate the role of NSUN2 in the context of DNA damage, we performed a Proximity Ligation Assay (PLA) using antibodies against NSUN2 and γH2AX (a DNA damage marker) following ionizing irradiation (IR)-induced DNA damage in Hela cells. Upon DNA damage induction (+IR), we detected an increased interaction between NSUN2 and γH2AX compared to the control condition (-IR) (Fig 1a).

**Fig. 1.**
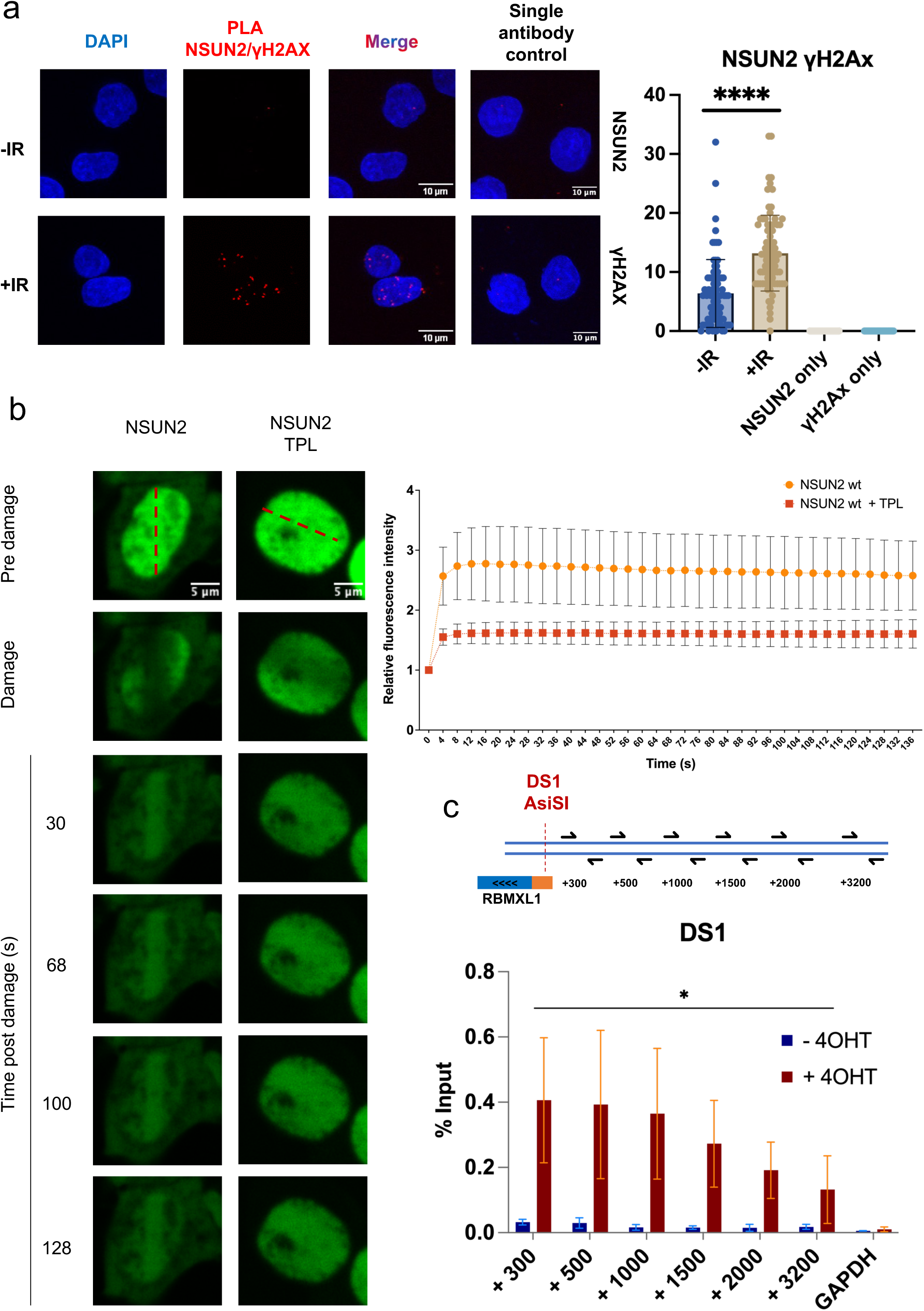
NSUN2 is recruited to double strand breaks. **a)** PLA of NSUN2 and γH2Ax with (+IR) or without (-IR) (10Gy, 15 min). As negative control single antibody assay was performed (NSUN2 and γH2Ax alone). Left: representative microscope images. Right: quantification of PLA nuclear foci. Errors bars, mean ± SD. N>100. Statistical significance was determined using non-parametric Mann-Whitney test. ****p ≤ 0.0001. **b)** Laser induced DNA damage of HEK 293T cells transiently transfected with Neon-Green-NSUN2 plasmid and incubation with or without Triptolide (TPL, 1h 10μM). Left: Representative confocal microscopy images before and after the laser induced damage. Right: quantification of relative Neon-Green signal (background signal/ROI signal) (n>15) along the time considered (0-132 s). Error bars: mean ± SEM. Statistical significance was determined by two-way ANOVA with multiple comparison test (p ≤ 0.0001). **c)** Top panel: Drawing depicting the *Asi*SI site DS1 (red dashed line), position of qRT-PCR primers, distance from target site is represented in nucleotides. Bottom panel: ChIP analysis of NSUN2 occupancy at DS1 and GAPDH sites. (C–F). Statistical significance was determined using non-parametric Mann-Whitney test. Error bars: mean ± SEM, *n* = 3. *p ≤ 0.05.

Furthermore, to assess the dynamics of NSUN2 recruitment to DNA damage sites, we employed live-cell microscopy combined with laser micro-irradiation. Initially, we transiently transfected HEK293T cells with Neon-Green-tagged NSUN2 wt and treated the cells with the RNAPII transcription inhibitor Triptolide (TPL). NSUN2 wt was rapidly recruited to DSBs, and its accumulation at laser stripes was reduced by TPL (Fig. 1b). To determine whether its recruitment to DSBs is dependent on its methylation activity, we subjected cells transfected with Neon-Green-NSUN2 wt and the Neon-Green-NSUN2 K190M mutant to laser micro-irradiation and observed that the recruitment kinetics were not significantly different between the wt and the K190M mutant. However, transcription inhibition by TPL reduced the recruitment of NSUN2 K190M to the laser stripes (Extended Data Fig. 1).

Additionally, to confirm the NSUN2 recruitment to DSBs, we employed the DiVA system (AsiSI-RE-U2OS), where an ER-fused AsiSI restriction enzyme translocates to the nucleus upon tamoxifen induction (+4OHT) to cleave DNA at specific sites, simulating DSBs. This cellular system is commonly used to study DSBs in a sequence specific manner ^26, 27^. We performed NSUN2 Chromatin immunoprecipitation (ChIP) on DSB induced in the transcriptionally active gene RBMXL1 (DS1) and at non-DSB exonic site (GAPDH). Following the induction of AsiSI-mediated DSB, we observed an enrichment of NSUN2 at DS1 and this enrichment decreased with the distance from the DS1 site. We did not detect NSUN2 at the control GAPDH locus (Fig. 1c).

These data suggest that the NSUN2 protein is present at DSBs. The recruitment of NSUN2 to the DNA damage site is independent of its RNA methylation activity, but requires active RNAPII transcription.

### NSUN2 binds and methylates newly synthesized antisense DNA damage-responsive transcripts (DARTs)

Giver that NSUN2’s recruitment to DSB depends on active RNAPII transcription and that NSUN2 is an RNA methyltransferase, we hypothesized that it might interact with RNA transcripts derived from DSBs. To investigate the interaction between NSUN2 and DARTs, we developed an advanced, modified version of FISH-PLA method termed zC-FISH-PLA. We designed specific DNA probes to target a 1 kb region surrounding DS2, which hybridize with either antisense (DS2 AS) or sense (DS2 SS) DARTs. Additionally, we took advantage of the distinctive property of the 5-azacytidine (zC) analogue, which incorporates into RNA molecules, forming a stable, covalent and irreversible bond between NSUN2 and the RNA. This leads to the zC-RNA-NSUN2 complex formation, signifying the active addition of m5C to RNA transcripts (Fig. 2a). Following DSBs induction, we observed that NSUN2 protein interacted with antisense DARTs (DS2 AS) but not with sense DARTs (DS2 SS) (Fig. 2b and Extended Data Fig. 2a). We also investigated the formation of zC-FISH-PLA nuclear foci under various control conditions. The presence of TPL reduced the number of NSUN2/RNA foci, verifying the zC-FISH-PLA specificity (Fig. 2b). Additionally, the depletion of NSUN2 (using siNSUN2) significantly decreased the number of detectable NSUN2/RNA foci (Fig. 2b). Utilizing scrambled DNA probes, another negative control, resulted in no formation of NSUN2/RNA foci (Fig. 2b). Collectively, these data imply that our zC-FISH-PLA is a highly specific and novel approach for detecting the interaction between NSUN2 and methylated RNA. Furthermore, zC-FISH-PLA did not show any detectable NSUN2/RNA foci when we used probes specific to an intergenic AsiSI site, DS3 (Extended Data Fig. 2b). Since the DNMT2 protein has been observed to methylate R-loop structures after H_2_O_2_-induced DNA damage, we also examined the interaction between DNMT2 and DS2 AS and DS2 SS DARTs using zC-FISH-PLA, but did not detect any foci (Extended Data Fig. 2c and d).

**Fig. 2.**
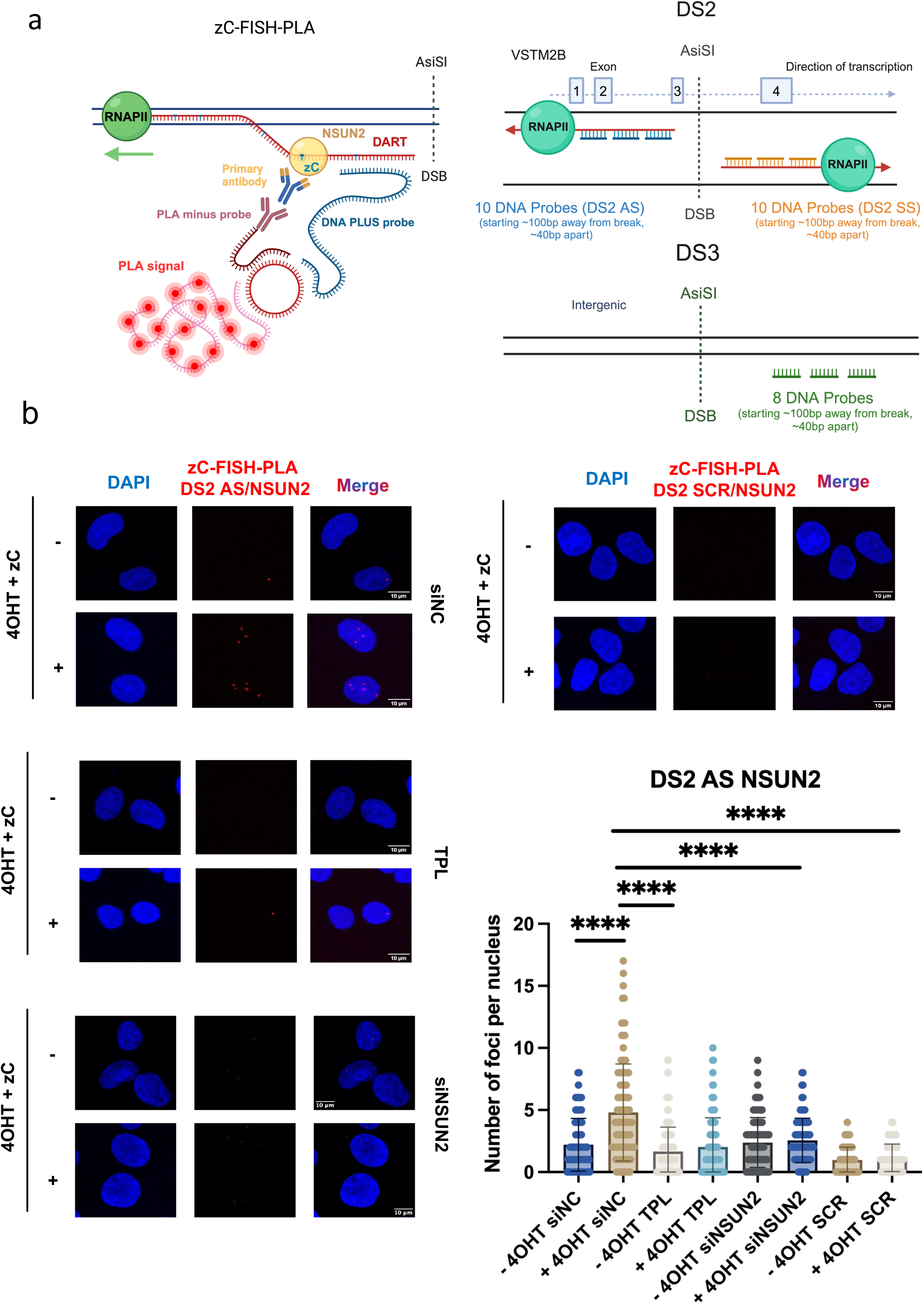
NSUN2 binds to and methylates antisense DNA Associated RNA Transcripts (DARTs) **a)** Illustration of zC-FISH-PLA assay: 5-Aza-Cytidine analogues (zC) are incorporated into the antisense DARTs and trap NSUN2 protein on the RNA molecule. Incubation with primary antibody against NSUN2 and DNA probes complementary to antisense RNA, DNA PLA minus probe and secondary PLA plus probe are ligated acting as a primer for rolling-circle amplification. Fluorescently-labelled oligos then hybridize to the complementary sequences within the amplicon producing discrete foci (left). Structure of *Asi*SI sites DS2 (VSTM2B) and intergenic DS3, antisense and sense DNA probes (right). **b)** Representative microscope images showing zC-FISH-PLA of NSUN2 and DNA probes (DS2 AS and DS2 SCR) with (+4OHT) or without (-4OHT) and (Z)-4-Hydroxytamoxifen incubation for 4h in NSUN2 depleted (siNSUN2), siNegative Control (siNC), and Triptolide treatment (TPL, 1h, 10μM). Quantification plot zC-FISH-PLA nuclear foci. Errors bars, mean ± SD. Statistical significance was determined using non-parametric Mann-Whitney test. ****p ≤ 0.0001.

Our thorough and targeted zC-FISH-PLA data support the hypothesis that NSUN2, as opposed to DNMT2, selectively binds to and methylates the newly synthesized antisense DARTs.

### Depletion of NSUN2 increases levels of nascent antisense RNA transcript levels at DSBs

Previous studies have indicated that the m5C modification deposited by NSUN2 contribute to increased stability of the mRNA molecules^28, 29^. To investigate whether NSUN2 plays a role in the regulation of the stability of DARTs, we performed Chromatin-bound RNA sequencing (Chr-RNA-seq) with strand-specific analysis of nascent RNA transcripts in wt (siNC) and NSUN2 knockdown (siNSUN2) cells after inducing AsiSI-derived DSBs (+4OHT). Principal Component Analysis (PCA) validated the high consistency among Chr-RNA-seq samples (Extended Data Fig. 3a). DSBs (as cut sites) were identified and mapped by BLESS technique^30^. Heatmap analysis of Chr-RNA-seq data revealed that NSUN2 depletion resulted to a significant increase in antisense DART levels (Fig. 3a). Further comparison of nascent antisense and sense DARTs derived from all AsiSI sites, confirmed that antisense DARTs levels were significantly higher that sense transcript levels following NSUN2 depletion (Fig. 3b, c, d, e and Extended data Fig. 3b and c).

**Fig. 3.**
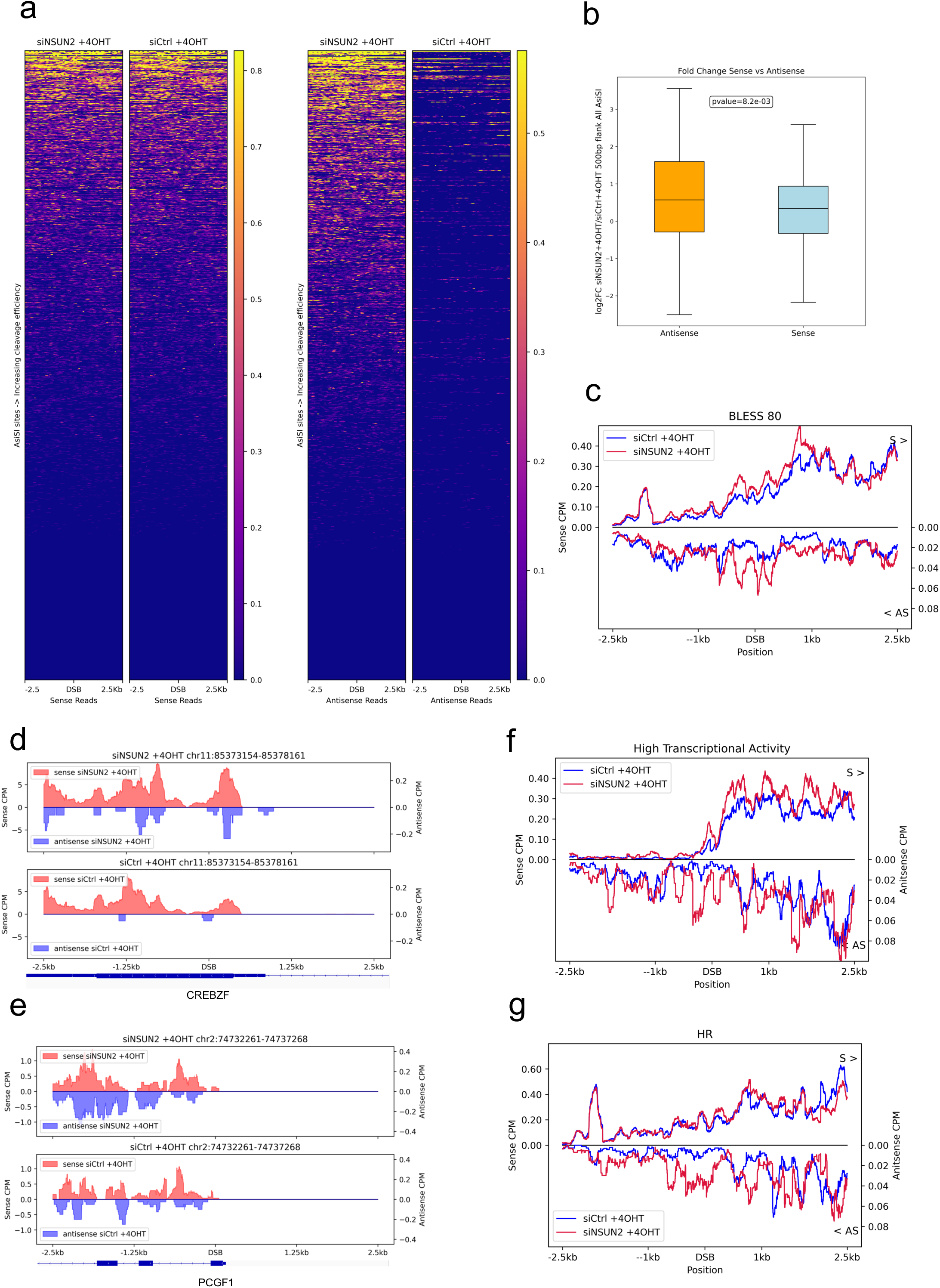
NSUN2 depletion leads to stabilization of antisense DARTs. **a)** Heatmaps show sense (left) and antisense (right) nascent RNA (chrRNA-seq) read coverage across all annotated AsiSI sites sorted based on cleavage efficiency in both NSUN2 knockdown and control cells with damage induction (+4-OHT). **b)** Box plots show log2FoldChange of chrRNA-Seq coverage of sense reads and antisense reads upon NSUN2 knockdown in damage conditions compared to control for all AsiSI (+/- 500bp). Wilcoxon 2 sample test is used for statistical testing of medians between sense and antisense log2FoldChange distribution. **c)** Metagene plot shows chrRNA-Seq sense and antisense coverage in control (siCtrl) and NSUN2 knockdown cells with damage induction (+4-OHT) around 2.5kb flank region of BLESS 80 AsiSI sites (n=80). The reference genome is human hg19. **d-e)** Representative snapshots of individual genes showing sense and antisense chrRNA-Seq coverage in NSUN2 knockdown and control with damage induction around 2.5kb flank region of AsiSI cut. The specific loci information is listed on top of the snapshots. The reference genome is human hg19. **f)** Metagene plot shows chrRNA-Seq sense and antisense coverage in control (siCtrl) and NSUN2 knockdown cells with damage induction (+4-OHT) around 2.5kb flank region of AsiSI sites associated with highly transcribed genes. The reference genome is human hg19. **g)** Metagene plot shows chrRNA-Seq sense and antisense coverage in control (siCtrl) and NSUN2 knockdown cells with damage induction (+4-OHT) around 2.5kb flank region of AsiSI sites which are HR prone. The reference genome is human hg19.

Additionally, metagene analysis of BLESS 80 AsiSI sites (all cut sites) showed a considerable up-regulation of antisense DARTs and a slight increased in sense transcripts within a 1 kb range of DSBs (Fig. 3c and Extended data Fig. 3b-d). To further investigate the impact of NSUN2 depletion on nascent antisense and sense transcript levels, we categorized DSBs based on their proximity to regions with either high or low transcriptional activity. NSUN2 depletion led to a significant increase in both sense and antisense DART levels at DSBs close to actively transcribing genes, whereas in transcriptionally repressed regions, only antisense DART levels were elevated in the absence of NSUN2 (Fig. 3f and Extended data Fig.4a-c).

Furthermore, AsiSI cut sites in the DIvA system can be classified into HR or NHEJ prone DSBs based on correlation ratio using ChIP-Seq data coverage of RAD51, well known facilitator of HR, (AsiSI site +-4kb) to XRCC4, known NHEJ factor, coverage (AsiSI site +-1kb). The top 30 sites were annotated as HR-prone due to higher RAD51 coverage, and the bottom 30 as NHEJ-prone due to a higher XRCC4 ratio^30^. NSUN2 depletion significantly affected both sense and antisense DART levels at HR-prone sites, while at NHEJ-prone DSBs, only antisense DARTs were significantly increased (Fig. 3g and Extended data Fig.4d-f).

These data imply that NSUN2 may regulate RNAPII transcription or possibly the processing of DARTs. Observing that the increased levels of DARTs were confined to specific regions in close proximity to DSBs, we further investigated the role of NSUN2 in the processing of DARTs.

### NSUN2 interacts with DICER

NSUN2 does not exhibit an RNA cleavage activity. Therefore, we hypothesized that its influence on the processing of DARTs might be indirect. RNA processing enzymes, such as DROSHA and DICER are known to localize near DSBs. Our previous work has suggested a role for DICER in DNA repair^9–12^.

Consequently, we hypothesized that NSUN2 might interact with DICER. To investigate this, we utilized Surface plasmon resonance (SPR) with purified NSUN2 and DICER proteins (Extended Data Fig. 5a and b). We conducted the assay using a serial DICER dilution (as the analyte) ranging from 0.6μM to 1.2nM to determine its interaction with NSUN2 (serving as the ligand). A distinct peak at 25 seconds confirmed the *in vitro* interaction between these two proteins (Fig. 4a).

**Fig. 4.**
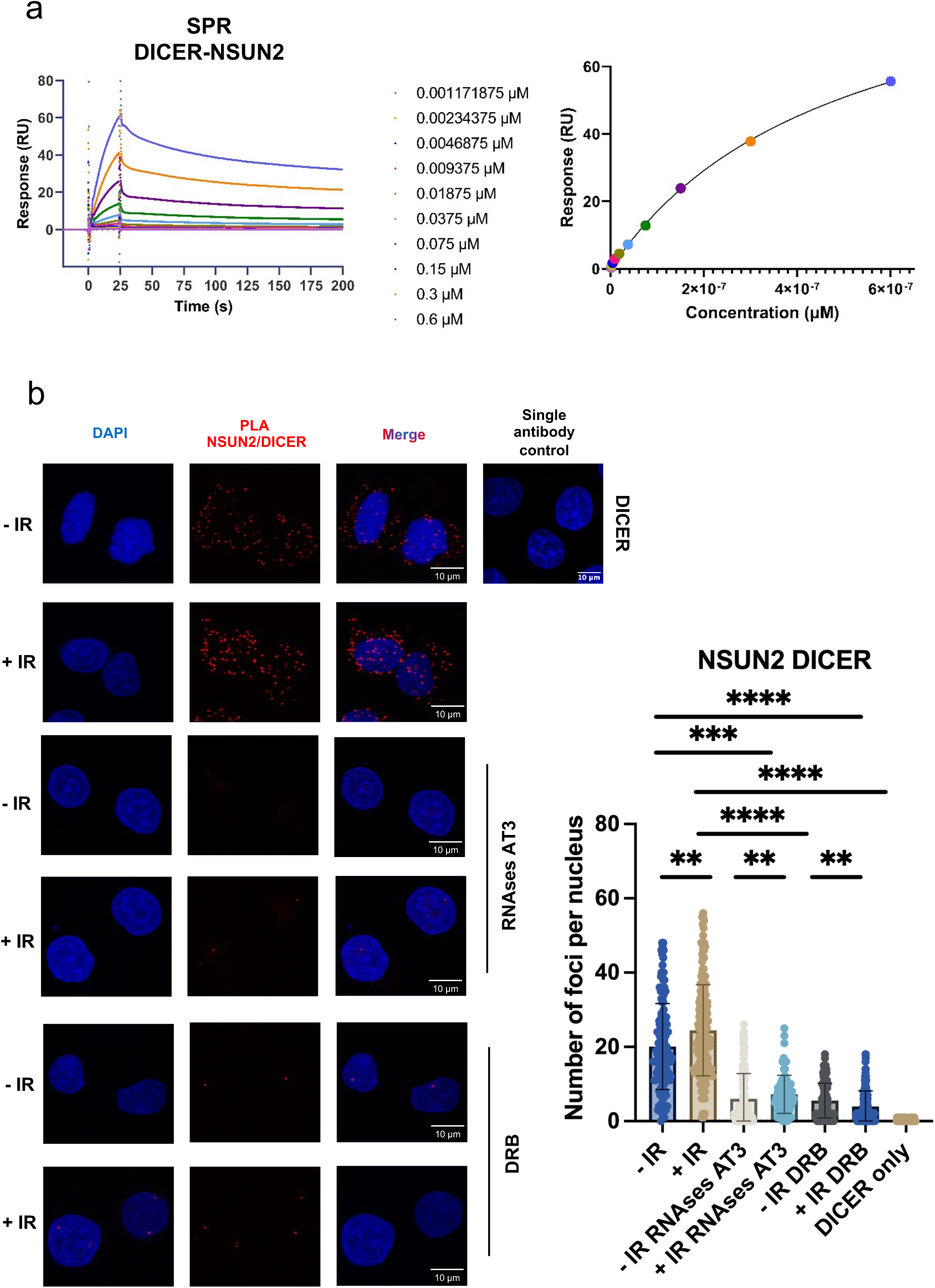
NSUN2 interacts with DICER. **a)** Left: Sensogram of showing signals for DICER, injected at various concentrations (0.6 *μ*M-1.2nM), binding to NSUN2 captured on the sensor surface. Right: curve dose-response of NSUN2-DICER association (Response versus Concentration), *K_d_* ≅0.5 *μ*M. **b)** PLA of NSUN2 and DICER with (+IR) or without (-IR) Ionizing Radiation (10Gy, 15 min), post PFA fixation treatment with RNAse A (100 *μ*g), RNAse T1(1000U) and RNAse Short Cut III (2000U) for 2h at room temperature (AT3) or in presence of transcription inhibitor, DRB, (100 *μ*M 2h). As a negative control single antibody assay was performed (DICER only). Left: representative microscope images. Right: quantification of PLA nuclear foci. Errors bars, mean ± SD. N>100. Statistical significance was determined using non-parametric Mann-Whitney test. ****p ≤ 0.0001, \*\*\**p* ≤ 0.001 and \*\**p* ≤ 0.01.

To test whether NSUN2 interacts with DICER in living cells, we performed a PLA assay in irradiated HeLa cells. We identified numerous NSUN2-DICER PLA foci within both the nucleus and cytoplasmatic compartments. Interestingly, IR treatment triggered an increase in the number of nuclear NSUN2-DICER PLA foci (Fig. 4b). To further dissect the NSUN2-DICER interaction, we applied a post-fixation treatment with various RNAses (*i.e.* RNAse A, RNAse T1 and RNAse III (AT3)) and detected a significant reduction of NSUN2-DICER PLA foci under both undamage and damage conditions (-IR and +IR, respectively). A significant reduction in NSUN2-DICER PLA foci number was also observed in the presence of the transcription inhibitor DRB, indicating that the interaction between NSUN2 and DICER is transcription-dependent (Fig. 4b). Finally, to test the potential interaction between DNMT2 and DICER, we employed DNMT2-DICER PLA, but no DNMT2-DICER PLA nuclear foci were detected (Extended Data Fig. 5c).

These data indicate that DICER specifically interacts with NSUN2 in an RNA-dependent manner.

### DICER binds to newly synthesized antisense DARTs

Our previous work demonstrated that DICER is recruited to AsiSI-induced DSBs and this recruitment is RNA-dependent^12^. To test the hypothesis that DICER is recruited to DARTs, we performed a FISH-PLA assay using DNA probes specific to DS2 AS in conjunction with a DICER antibody. Upon DSB induction, we observed an increased number of DS2 AS-DICER FISH-PLA foci, which suggests that DICER indeed binds to the antisense DARTs (Fig. 5a). Interestingly, NSUN2 depletion resulted in a significant increase in FISH-PLA foci compared to the negative control (siNC) after induction of DSBs. These data align with the observed increase in DARTs following NSUN2 depletion. Neither the depletion of DICER nor the use of scrambled DNA probes (SCR DS2) resulted in the formation of FISH-PLA foci. Furthermore, we conducted a comparative analysis of metagene profiles of DARTs with DICER ChIP-seq data^12^. This comparison revealed a positive correlation between DICER occupancy and the elevated levels of antisense DARTs at BLESS 80 AsiSI sites in the absence of NSUN2 (Fig. 5b).

**Fig. 5.**
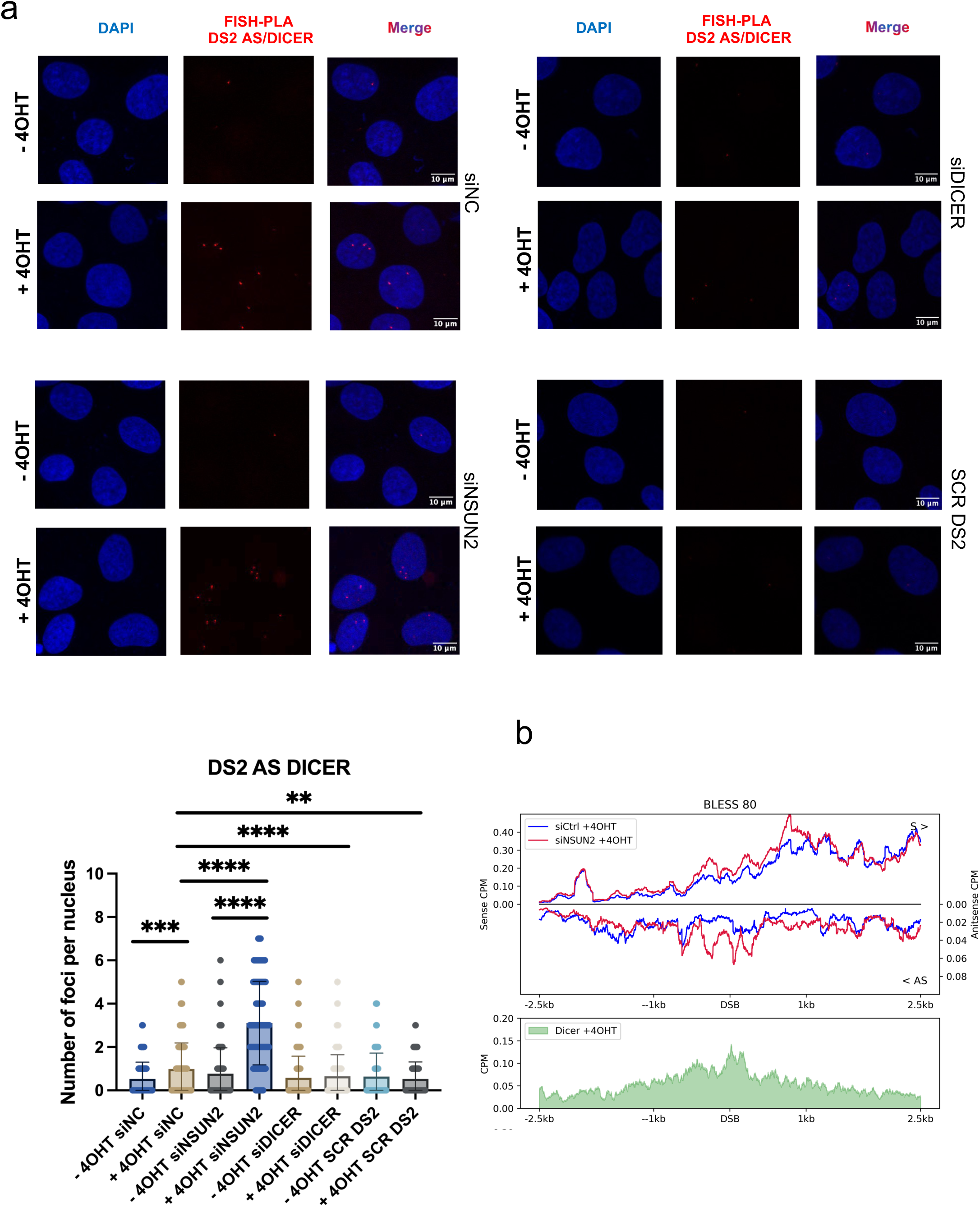
DICER binds to *de novo* antisense DARTs. **a)** Representative microscope images of FISH-PLA of DICER and DNA probes (DS2 AS and DS2 SCR) with (+4OHT) or without (-4OHT) and (Z)-4-Hydroxytamoxifen incubation for 4h in NSUN2 depleted (siNSUN2), siNegative Control (siNC), and Triptolide treatment (TPL, 1h, 10μM). Quantification plot zC-FISH-PLA nuclear foci. Errors bars, mean ± SD. Statistical significance was determined using non-parametric Mann-Whitney test. ****p ≤ 0.0001, \*\*\**p* ≤ 0.001 and \*\**p* ≤ 0.01. **b)** Metagene plot shows chrRNA-Seq sense and antisense coverage in control (siCtrl) and NSUN2 knockdown cells with damage induction (+4-OHT) around 2.5kb flank region of BLESS 80 AsiSI sites (n=80). The reference genome is human hg19. DICER ChIP-seq^12^ metagene plot of BLESS 80 *Asi*Si sites after (Z)-4-Hydroxytamoxifen incubation (4h).

These data indicate that DICER is actively recruited to antisense DARTs and that the absence of NSUN2 protein results in an increased interaction between DICER and DARTs.

### NSUN2 facilitates DICER processing of DART-associated R-loops

Our results on NSUN2 and DICER interaction and their collective binding to antisense DARTs, along with the observation of increased DART levels under NSUN2 knockdown conditions, led us to hypothesize that NSUN2 may aid in the DICER-mediated processing of DARTs. To explore this hypothesis, we purified recombinant NSUN2 wt, NSUN2 K190M (non-methylating mutant), DICER wt and DICER DEDE (a catalytically inactive mutant) proteins (Extended data Fig. 5a and b). Additionally, we synthesized various RNA substrates: single-stranded RNA (ssRNA), double-stranded RNA (dsRNA), DNA:RNA heteroduplex, R-loop and an R-loop with a 5’-tail (Fig. 6a, b and Extended Data Fig. 7a-c). All RNA sequences were *in vitro* transcribed (IVT) and contained the sequence of the antisense DS2, which we used in *in vivo* experiments (400 nt from the AsiSI site). The R-loop + 5’-tail consists of DS2 400 nt and 120 nt scrambled sequence.

**Fig. 6.**
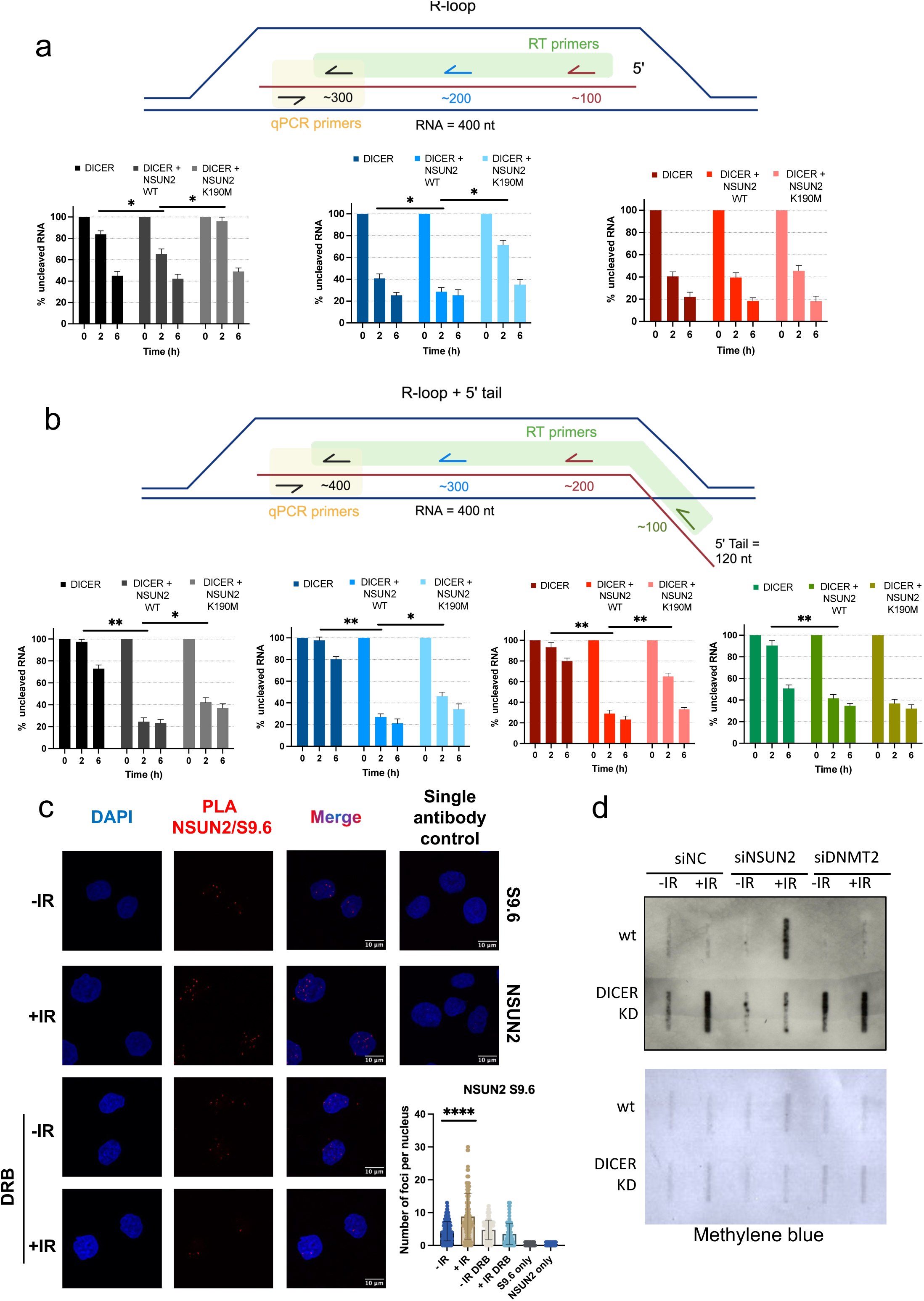
NSUN2 promotes DICER processing of DART-associated R-loops. **a)** Top: Schematic representation of *in vitro* transcribed (IVT) DS2 antisense RNA forming R-loop with RT primers: (red arrow ∼100 nt, blue arrow ∼200 nt and black arrow ∼300nt away from the *Asi*Si site, highlighted in green), and qPCR primers (black arrows, highlighted in yellow). Bottom: quantification plots of cleaved RNA at 0, 2h and 6h time points in presence of DICER only, DICER + NSUN2 wt and DICER + NSUN2 K190M. Black, blue and red bars correspond to black, blue and red primer specific RT, respectively. Errors bars, mean ± SD. N=2. Statistical significance was determined using non-parametric Mann-Whitney test, \**p* ≤ 0.05. **b)** Top: Schematic representation of IVT DS2 antisense RNA forming R-loop 5’-tail with RT primers: (green arrow ∼100 nt, red arrow ∼200 nt, blue arrow ∼300nt and black arrow ∼300nt away from the *Asi*Si site, highlighted in green), and qPCR primers (black arrows, highlighted in yellow). Bottom: quantification plots of cleaved RNA at 0, 2h and 6h time points in presence of DICER only, DICER + NSUN2 wt and DICER + NSUN2 K190M. Black, blue, red and green bars correspond to black, blue, red and green primer specific RT, respectively. Errors bars, mean ± SD. N=2. Statistical significance was determined using non-parametric Mann-Whitney test. \*\**p* ≤ 0.01 and \**p* ≤ 0.05. **c)** PLA of NSUN2 and S9.6 with (+IR) or without (-IR) Ionizing Radiation (10Gy, 15 min), or in presence of transcription inhibitor, DRB (100 *μ*M 2h). As a negative control single antibody assay was performed (S9.6 and NSUN2 only). Top and left: representative microscope images. Bottom: quantification of PLA nuclear foci. Errors bars, mean ± SD. N>100. Statistical significance was determined using non-parametric Mann-Whitney test. ****p ≤ 0.0001. **d)** Top: Slot Blot probing with m5C antibody on RNA isolated from cells treated with (+IR) or without (-IR) Ionizing Radiation (10Gy, 10 min), in NSUN2 depleted (siNSUN2), DNMT2 depleted (siDNMT2), siNegative Control (siNC), and HEK 293T or HEK 2B2 depleted for DICER protein. Bottom: Methylene blue blot used as a loading control.

Initially, we tested the cleavage of ssRNA and dsRNA by different combinations of NSUN2 and DICER mutants *in vitro* followed by denaturing urea PAGE. The NSUN2 wt alone showed no RNA processing, confirming its lack of RNA cleavage activity. However, when NSUN2 wt was combined with DICER wt, there was significant cleavage of both ssRNA and dsRNA substrates in a time-dependent manner. No significant processing of ssRNA and dsRNA species was detected with NSUN2 wt and mutant the DICER DEDE mutant. Interestingly, combining NSUN2 K190M with DICER wild type led to ssRNA or dsRNA cleavage comparable to that of NSUN2 wild type with DICER wild type, indicating that m5C deposition by NSUN2 does not significantly affect DICER’s cleavage of these RNA substrates. (Extended Data Fig. 6a and b).

To further understand the role of NSUN2 during DART processing by DICER, we utilized a quantitative method, RT-qPCR, to measure the RNA cleavage rate post *in vitro* reaction. An equal amounts of *in vitro* RNA cleavage reaction were collected at different time points (*i.e.* 0, 2h, 6h), reverse transcribed with specific RT primers hybridizing to the 3’-end (black), middle part (blue), 5’-end (red) and 5’-end tail (green) of the RNA substrates and followed by quantitative real time qPCR. Assessment of the DICER wt cleavage rate of ssRNA, dsRNA substrates by RT-qPCR confirmed robust RNA processing, which was not affected by the presence of either NSUN2 wt or K190 mutant. These data were consistent with our previous cleavage assay results (Extended data Fig. 6a and b), suggesting that the RT-qPCR is a reliable method to determine the DICER cleavage rate of RNA species (Extended date Fig. 7a and b). Interestingly, DICER wt, whether alone or in combination with NSUN2 wt or K190 mutant, did not cleave DNA:RNA hybrid substrates (Extended date Fig. 7c).

**Fig. 7.**
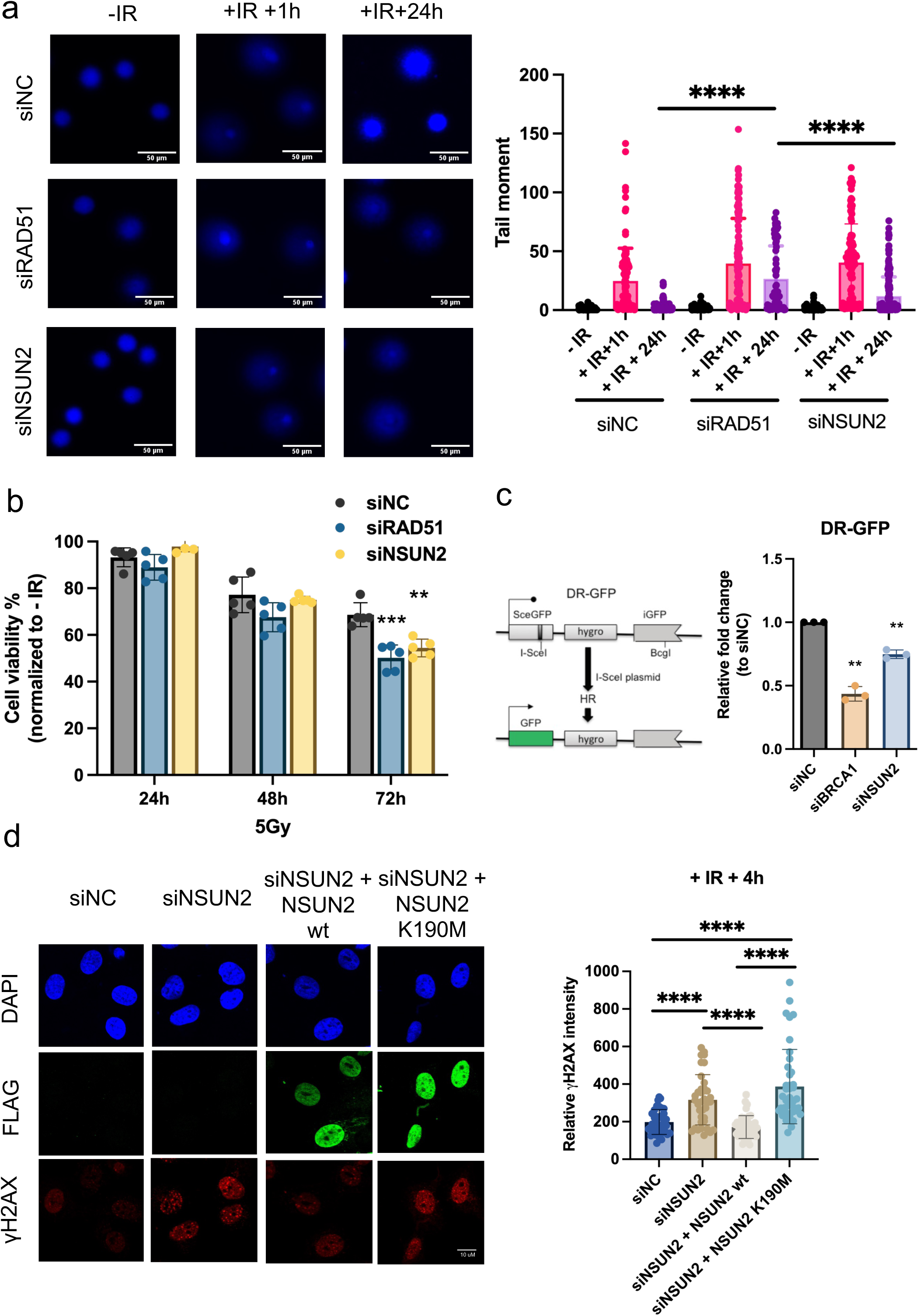
Depletion of NSUN2 delays DNA repair by HR. **a)** Left: Representative images of Comet assay with (+IR +1h, +IR 24h,) or without (-IR) Ionizing Radiation (5Gy, 15 min), in NSUN2 depleted (siNSUN2), RAD51 depleted (siRAD51), siNegative Control (siNC). Right: Quantification plot of the tail moment. Errors bars, mean ± SD. Statistical significance was determined using non-parametric Mann-Whitney test. ****p ≤ 0.0001. **b)** MTS assay of HeLa cells irradiated with 5 Gy at 24h, 48h and 72h in NSUN2 depleted (siNSUN2), RAD51 depleted (siRAD51), siNegative Control (siNC) conditions. **c)** Schematic illustration of DR-GFP system and efficiency of HR repair in NSUN2 depleted (siNSUN2), BRCA1 depleted (siBRCA1), siNegative Control (siNC) conditions. Errors bars, mean ± SD. Statistical significance was determined using non-parametric Mann-Whitney test. \*\**p* ≤ 0.01. **d)** Left: representative immunofluorescence images of FLAG-tagged NSUN2 wt, FLAG-tagged NSUN2 K190M and γH2Ax in NSUN2 depleted (siNSUN2), siNegative Control (siNC), NSUN2 wt rescued (siNSUN2 + NSUN2 wt) and NSUN2 K190M rescued (siNSUN2 + NSUN2 K190M) irradiated with 5 Gy at 4h. Right: Quantification plot of the γH2Ax intensity. Errors bars, mean ± SD. Statistical significance was determined using non-parametric Mann-Whitney test. ****p ≤ 0.0001.

Previous studies showed that DICER is able to process non-canonical RNA substrates such as tRNAs^31^ or R-loops^32^. R-loops are known to be associated with DSBs and are formed by DARTs hybridization to DNA templates^8, 16, 33–38^. Therefore, we also performed an *in vitro* DICER cleavage assay using an R-loop substrate. We detected DICER mediated and time-dependent cleavage of the R-loop with all three primer sets, although this was lot less efficient than DICER cleavage of ss or dsRNA. This processing was increased by the presence of NSUN2 wt and significantly reduced by NSUN2 K190 mutant for two of three tested primer sets (Fig. 6a).

Considering that R-loops in cells would include an ssRNA tail due to nascent RNA hybridization to the DNA template, we tested DICER’s processing of the R-loop with a 5’-tail substrate. Interestingly, we observed only a weak DICER processing of the R-loop 5’-end tail substrate. However, DICER was robustly stimulated by the presence of NSUN2 wt and significantly reduced by NSUN2 K190M for all three primer sets probing within the R-loop substrate (Fig. 6b). We also noticed that the lack of NSUN2-mediated m5C deposition on RNA did not affects DICER processing next to the junction region where the RNA undergoes local conformational transition from ssRNA to R-loop structure (green primer). Given that the presence of m5C modification has a stabilization effect on both DNA and RNA thermodynamic properties, we reasoned that NSUN2-dependent m5C deposition may assist DICER processing modulating the local conformational geometry toward a more favourable structure for DICER interaction and cleavage (Fig. 6b).

Next, we investigated whether NSUN2 can be detected in proximity to R-loops *in vivo*. We performed a NSUN2-S9.6 optimised PLA assay^39^ and detected an increased number of nuclear NSUN2-S9.6 foci in DNA damage conditions, which was reduced in the presence of transcription inhibitor DRB (Fig. 6c). Finally, we isolated nuclear RNA and performed a slot blot using m5C antibody. A comparison of the m5C signal in HEK wt cells depleted of either DNMT2 or NSUN2, revealed a strong m5C signal originated from DNMT2 rather than NSUN2 activity (Fig. 6d). Furthermore, depletion of DICER resulted in a strong accumulation of m5C-modified nuclear RNA molecules upon DNA damage, suggesting that indeed DICER cleaves methylated nuclear RNA. Depletion of NSUN2, but not of DNMT2, dramatically reduced the m5C signal on nuclear RNA in damage conditions, suggesting that DNMT2 is able to modify RNA molecules, which are not DICER substrates (Fig. 6d).

Taken together these data imply that NSUN2, but not DNMT2, chaperones DICER in the processing of the DART-associated R-loops.

### NSUN2 methylation activity is crucial for efficient DNA damage repair

To investigate the role of NSUN2 is DSBs repair *in vivo*, we performed a direct comet assay in HeLa cells. Quantification of the tail moment in wt condition revealed presence of DNA damage at 1h time point, which were nearly completely repaired by 24 hours post irradiation. However, both RAD51 and NSUN2 depleted cells revealed a persistent DSBs even at 24h time point (Fig. 7a and Extended Data Fig. 8a). Furthermore, depletion of NSUN2 had a significant effect on cellular viability following irradiation (Fig. 7b). To test, which DNA damage repair pathway is affected by NSUN2 depletion, we used plasmid-based GFP-reporters DR (homologous recombination) and EJ5 (non-homologous end joining) in wt and NSUN2 KD conditions. NSUN2 depletion resulted in impaired homologous recombination repair efficiency (DR) while no significant changes have been detected in repair by non-homologous end joining (Fig. 7c and Extended Data Fig. 8b and c; DNA-PK inhibitor Wortmannin and depletion of BRCA1 were used as negative controls).

To investigate whether NSUN2 itself or its methylation activity are important for the DSB repair, we analysed the γH2AX clearance in wt and NSUN2 depleted cells and rescued with FLAG-NSUN2 wt and FLAG-NSUN2 K190M conditions at 1, 4 and 24 hour time points. Depletion of NSUN2 protein had a significant impact on the clearance of γH2AX at 4h and 24h post IR. This repair defect was rescued by addition of NSUN2 wt, but not by NSUN2 K190M mutant (Fig. 7d and Extended data Fig. 8d).

These results suggest that NSUN2 and its methylation activity are crucial for the efficient repair of DSBs.

## Discussion

Non-coding transcripts that originate from DSBs, such as DARTs are required for effective DNA repair ^7, 8^. These transcripts require synthesis by RNAPII, as well as further modifications, processing and eventual clearance. Our study reveals an unexpected role for the m5C methyltransferase NSUN2 in the context of DNA damage. NSUN2 is rapidly recruited to DSBs in transcription-dependent manner. Moreover, depletion of NSUN2 results in the stabilization of DARTs, presumably due to inefficient cleavage by DICER in the context of damage-associated R-loops.

A recent report indicated that DICER’s ability to cleave R-loop structures is minimal compared to its effectiveness on hairpin RNAs when tested *in vitro*^32^. Our *in vitro* RNA cleavage assays demonstrate that NSUN2 significantly stimulates DICER’s efficiency in processing non-canonical substrates, such as R-loop with 5’-tail. This improved activity of DICER depends on both a direct protein-protein interaction with NSUN2 and the presence of m5C modifications. The identification of NSUN2’s role in facilitating DICER’s cleavage of R-loop with a 5’ tail is particularly crucial, because this structure mimics the natural R-loop structures that consists of a DNA:RNA heteroduplex, a displaced ssDNA and a 5’ trailing ssRNA. Thus, the effective processing of R-loop with a 5’-tail by DICER in the presence of NSUN2 suggests their vital role in the clearance of DARTs. Given that NSUN2 methylates a variety of RNA substrates, it is conceivable that these modifications could facilitate DICER’s cleavage of other non-canonical RNA molecules.

NSUN2’s ability to augment DICER’s cleavage efficiency is not unique. Previous studies have identified protein partners of DICER, such as ADAR, an enzyme that converts Adenosine to Inosine, which also directly binds to DICER and enhances miRNA processing^40^. This raises the possibility that DICER might interact with various RNA modifiers, including both ADAR and NSUN2, influencing RNA metabolism through post-transcriptional modifications. Similar to ADAR, NSUN2 could preferentially enhance the cleavage of certain RNA species in response to specific signals.

Previous studies on m5C modifications following ROS-induced DNA damage has proposed that DNMT2, rather than NSUN2, was the primary methyltransferease for m5C deposition at DSBs^41^. It is crucial to note, however, that the photoactivable ROS generators used in those studies can cause a broad range of DNA damage, including single-strand breaks, DSBs, and oxidative lesions such as 8-oxoG. Another study indicated that DNMT2 is involved in resolution of oxidative stress^42^. In contrast, our study utilized the DIvA system that precisely induces DSBs^30^ to assess RNA m5C deposition during the DNA damage response (DDR). Our findings imply that m5C modifications on DARTs are indeed dependent on NSUN2, not DNMT2, emphasizing the need for accurate experimental models to study specific DNA damage types and their related RNA modifications.

Upon evaluating m5C levels on nuclear RNAs, we discovered a striking difference in the fate of RNAs methylated by NSUN2 compared to those modified by DNMT2 following DNA damage. NSUN2-methylated RNAs are quickly degraded by DICER post-damage induction, whereas DNMT2-methylated RNAs are not processed by DICER. This distinction underscores a unique regulatory pathway for RNA stability contingent on the specific methyltransferase responsible for m5C deposition.

In conclusion, our findings suggest a dual role for NSUN2 in the DNA damage response: firstly, through protein-protein interaction, NSUN2 assists DICER in the removal of DARTs; secondly, it influences the activity or localization of m5C reader proteins at DNA damage sites by depositing m5C on RNA. Overall, we propose that NSUN2 is an important factor in DNA repair.

## Materials and Methods

### Cell lines and cell culture

HeLa, HeLa EJ5-GFP, HeLa DR-GFP, HEK 293T, HEK 293T 2B2 (gift from the Filipowicz’s Lab), U2OS and AsiSI-ER U2OS (gift from Legube’s Lab) cells were maintained under exponential growth in high-glucose DMEM medium (Life Technologies #31966047) supplemented with 10% (vol/vol) heat inactivated fetal bovine serum (Sigma-Aldrich #F9665), 2 mM L-glutamine (Life Technologies #25030024) and 100 units/ml penicillin-streptomycin solution (Life Technologies, 15140122) at 37 °C with 5% CO_2_. Regular tests for mycoplasma contamination have been conducted by PCR assessment method. The DNA damage was generated by γ-rays source.

### Plasmid construction

ORF of human NSUN2, NSUN2 K190M, DICER DEDE has been obtained from IDT as double-stranded DNA fragments (*i.e.* gBlocks HiFi Gene Fragments). Production of FLAG-NSUN2, FLAG-NSUN2-K190M, NeonGreen-NSUN2, NeonGreen-NSUN2-K190M, pFastBac-DICER-DEDE, pFastBac-NSUN2 and pFastBac-NSUN2-K190M vectors has been obtained by Gibson assembly methodology and Gibson Assembly Master Mix (NEB # E2611S). DNA sequences of recombinant plasmids have been determined by Sanger sequencing.

### siRNA and plasmid transfection

siRNA molecules (60 nM) have been reverse-transfected using Lipofectamine RNAiMax (Life technologies #13778075) and following the manufacturers’ instructions. All experiments were conducted following a 48-hour period subsequent to siRNA transfections, unless otherwise indicated. siRNA sequences used are listed in the Supplemental information. Plasmid DNAs were forward-transfected Lipofectamine 3000 (Invitrogen, #L3000001) and following the manufacturers’ instructions.

### Proximity Ligation Assay (PLA)

PLA was performed by using Duolink In Situ Red Starter Kit Mouse/Rabbit (Merck, #DUO92101-1KT) according to the manufacturer’s instructions. Cells were seeded onto poly-L-lysine coated glass coverslip to reach 60% confluency before fixing with 4% paraformaldehyde (PFA) in PBS (Alfa Aesar # J61899) for 10 min at 37°C. Permeabilization step was performed with 0.1% Triton X-100 (Merck, #X100-100ML) in PBS for 10min at room temperature before blocking 1h at 37°C with Duolink Blocking buffer. Primary antibodies were diluted (1:1000) with Duolink Dilution buffer and incubated overnight at 4°C. Subsequent PLA probe incubation, ligation and amplification steps were performed following the manufacturer’s instructions. Then coverslips were mounted with DAPI solution (Sigma-Aldrich #DUO82040) and sealed with transparent nail varnish before imaging with Olympus FluoView Spectral FV1200 confocal microscope with 60X oil immersion objective. Images were processed with ImageJ software and quantified by using CellProfiler with Speckle Counting pipeline.

### Chromatin Immunoprecipitation (ChIP)

ChIP and qPCR were performed by standard procedures as previously described^8^, with some minor changes. After incubation with 400nM (Z)-4-Hydroxytamoxifen (4OHT, Sigma-Aldrich #H7904) for 4h, 1 × 10^7^ AsiSI-ER-U2OS cells were double crosslinked with 1.5 mM EGS (ethylene glycol bis (succinimidyl succinate)) (ThermoFisher #21565) for 20 min at room temperature followed by 1% Formaldehyde (Merck, 252549-100ML) incubation for 10 min at 37°C. Crosslinking reaction was then and quenched by freshly prepared Glycine solution to a final concentration of 125 mM for 10 min at 37°C. Following nuclei lysis, chromatin was sheared by Bioruptor Pico device (Diagenode) for 30 min (ultra-high power, 30s ON/OFF). The sheared chromatin was collected by centrifugation, diluted and precleared with 30 μL protein A/G agarose beads (Merck-Millipore, 16-157/16-201) for 1h at 4°C. 3μg of antibody was used to isolate protein-DNA complex overnight at 4°C on rotating wheel. The protein-DNA complex was then pulled down by adding 60 μL protein A/G agarose beads for 1h at 4°C. After washes, elution and treatment with RNAse A (ThermoScientific # EN0531) and Proteinase K (NEB # P8107S) overnight at 65°C, DNA was purified by Phenol/chloroform/isoamyl alcohol solution (25:24:1, v/v) (Invitrogen #15593031) and isopropanol precipitated. qPCR was performed in triplicate using SensiMix SYBR (Bioline #QT65005) and 10 µM each of forward and reverse primers (listed in Supplemental information) on Rotor-Gene Q (QIAGEN). The thermal cycling conditions included an initial denaturation step at 95°C for 10 min, followed by 45 cycles at 95°C for 15s, 62°C for 15s, and 72°C for 20s. Melting curve analysis was conducted at the end of the qPCR amplifications.

### FISH-PLA

FISH-PLA and zC-FISH-PLA assays were performed according to previous publication (Alagia et al. 2023), with some minor changes. For zC-FISH-PLA experiments cells were incubated with 5 *μ*M of 5-Azacytidine (zC, Sigma-Aldrich #A2385) for 1h at 37°C before incubation of 400nM (Z)-4-Hydroxytamoxifen (4OHT, Sigma-Aldrich #H7904) for 4h.

### Surface Plasmon Resonance (SPR)

SPR was performed using Biacore S200. Purified 6xHis-NSUN2, at 0.2 *μ*g/mL, was immobilised on CM5 chip at pH 4.0 by amine coupling. pH (4.0-5.5) was scooted for best immobilisation conditions beforehand. Affinity assays were run with 2-fold serial dilutions of purified 6xHis-DICER in SPR running buffer (50 mM TrisHCl (pH 8), 50 mM NaCl, 5mM MgCl2, 2 mM TCEP, 0.05% Tween20).

### ChrRNA-seq sample preparation

The AsiSI-ER-U2OS cells at 70% confluency were incubated with 400nM 4-OHT for 4h. Then, the chromatin RNA samples for sequencing were prepared according to previous publications^31,43^. RNA was extracted from the chromatin samples using TRIzol LS (Invitrogen # 10296028) and 1-Bromo-3-chloropropane (Sigma-Aldrich # B9673) extraction, followed by Monarch Total RNA Miniprep Kit (NEB) according to the manufacturer’s instructions, and eluted in DEPC treated H_2_O.

### Protein purification

Recombinant NSUN2, NSUN2 K190M, DICER and DICER DEDE were produced according to previous publication (Alagia et al, 2023). Briefly, Spodoptera frugiperda (Sf9) cells were infected with P1 baculovirus at MOI = 1, generated with either pFastBac1-NSUN2, or pFastBac1-NSUN2 K190M, or pFastBac1-hDICER (Addgene #89144), or pFastBac1-hDICER-DEDE and the Bac-to-Bac expression system (Gibco # 10359016). All steps of purification were carried out at 4°C. Cells were harvested 72 h post-infection and the lysate was loaded on pre-equilibrated Ni-NTA agarose (QIAGEN #30210). 6xHis-NSUN2, 6xHis-NSUN2-K190M, 6xHis-DICER, 6xHis-DICER-DEDE were then eluted with a gradient of Imidazole. Fractions were run on SDS–PAGE and stained with Coomassie blue. Active fractions were concentrated, and buffer was exchanged with a 100 kDa cut-off for DICER and 50 kDa for NSUN2 concentrator (CYTIVA Vivaspin) and stored in storage buffer (50 mM Tris– HCl, pH 8, 50 mM NaCl, 5 mM MgCl2, 0.2% [mg/ml] BSA, 2 mM TCEP, 20% glycerol, and 1x protease inhibitor) at −80°C.

### Synthesis of DS2-derived dsDNAs for R-loop and ssDNA for DNA:RNA heteroduplex formation

Genomic DNA from U2OS cells has been isolated with Phenol/chloroform/isoamyl alcohol (25:24:1, v/v) (Invitrogen #15593031), precipitated with isopropanol and resuspended in water. DsDNA for R-loop (1kb) has been PCR synthetized using primers listed in Supplemental information. ssDNA for DNA:RNA heteroduplex (400nt) has been prepared using phosphorylated primer and Lambda Exonuclease (NEB # M0262S). Length of DNA Fragments were determined by agarose gel electrophoresis and then purified with Monarch DNA Gel Extraction Kit (NEB # T1020L). DNA concentration has been assessed by Nanodrop.

### *In vitro* transcribed RNA

DICER RNA substrates have been obtained using PCR synthetized linear DNA bearing T7 promoter and HiScribe T7 High Yield RNA Synthesis Kit (NEB #E2040S) following manufacturers’ instructions. After overnight reaction at 37°C and 4U Turbo DNAse treatment 1h at 37°C (Invitrogen #AM2238), RNA molecules have been purified by TRIzol and Isopropanol precipitation. RNA has been resuspended in DEPC treated H_2_O and stored at - 80°C.

### *In vitro* DICER cleavage assay

Equimolar quantity of T7 in vitro transcribed (IVT) RNAs (DS2_AS, DS2_SS and DS2_AS_tail) and either ssDNA or dsDNA were annealed by boiling at 98°C for 3 min and then slowly cool down to room temperature over a period of no less than 3 hours. ssRNA, dsRNA, DNA:RNA heteroduplex, R-loop and R-loop 5’-tail substrates were pre-incubated with either 1 μg of recombinant NSUN2 or NSUN2 K190M in DICER reaction buffer (50 mM Tris–HCl, pH 8, 300 mM NaCl, 20 mM Hepes, 5 mM MgCl_2_, 5mM CaCl_2_, 0.5mM DTT, 5% glycerol, and 1x protease inhibitor) supplemented with 0.32 mM SAM for 1 h at 37°C, then 1 μg of either recombinant DICER or DICER DEDE were added to the reaction and incubated for 6 h at 37°C. Aliquots were collected at certain time points (0, 2h, 6h) and treated with 4U Turbo DNAse for 1h at 37°C followed by 0.8 U Proteinase K for 1h at at 37°C prior isopropanol precipitation and resuspension in DEPC-treated H_2_O. Then RNAs were reverse transcribed using several RT primers and SuperScript III Reverse Trascriptase (Invitrogen # 18080093) following manufacturers’ instruction for gene specific primer. Before Sybr Green (Bioline # QT650-05) real time qPCR, samples have been incubated with 5U of RNAse H (NEB # M0297S) for 1h at 37°C

### Northern Blot

T7 in vitro transcribed (IVT) RNA (DS2_AS) was dephosphorylated with Quick CIP (NEB # M0525S) for 30 minutes at 37°C, followed by heat-inactivation at 80°C for 2 minutes. Then, RNA was 5’-labelled with 20 U of T4 Polynucleotide Kinase (3’ phosphatase minus) (NEB # M0236S) and 50 pmol of (gamma-^32^P)ATP (Perkin Elmer) for 1h at 37°C. Radiolabelled DS2_AS and either ssRNA (DS2_SS) or ssDNA were annealed by boiling at 98°C for 3 min and then slowly cool down to room temperature over a period of no less than 3 hours. ssRNA, dsRNA, DNA:RNA heteroduplex, were pre-incubated with either 1 μg of recombinant NSUN2 or NSUN2 K190M in DICER reaction buffer (50 mM Tris–HCl, pH 8, 300 mM NaCl, 20 mM Hepes, 5 mM MgCl_2_, 5mM CaCl_2_, 0.5mM DTT, 5% glycerol, and 1x protease inhibitor) supplemented with 0.32 mM SAM for 1 h at 37°C, then 1 μg of either recombinant DICER or DICER DEDE were added to the reaction and incubated for 6 h at 37°C. Aliquots were collected at certain time points (0, 2h, 6h), mixed with Novex TBE–urea loading dye (Invitrogen #LC6876) and immediately frozen on dry ice. Samples were run on a gel containing 7M Urea and 4% acrylamide/bis-acrylamide (29:1) in 1x TBE at 450 V for 1h. Then, the gel was dried and the radiolabelled RNA was visualized by autoradiography.

### Slot Blot

HEK 293T and HEK 293T 2B2 cells were incubated with Doxycycline (1 μg/μl) for 3 days, and reverse transfected with siNC, siNSUN2 and siDNMT2 (60nM). Cells were DNA damaged with 10 Gy, incubated for 10 min at 37°C and nuclear RNA was obtained by cellular fractionation. Cells were incubated with Hypotonic Buffer (20mM Tris-HCl pH 7.4, 10mM NaCl, 3mM MgCl_2_, 0.1% NP-40 in DEPC-treated H_2_O) for 5 min on ice, centrifuged at 500g for 5 min at 4°C, and the isolated nuclei were incubated with Cell Lysis Buffer (50mM Tris-HCl pH7.4, 150mM NaCl, 3mM EDTA, 3mM EGTA, 1% Triton X-100, 0.1% SDS, 2mM DTT in DEPC-treated H_2_O) for 5 min on ice. Lysed nuclei were subsequently added to TRIzol LS solution and 1-Bromo-3-chloropropane (3:1 v/v) and incubated for 5 min at 55°C on a thermomixer. RNA was then isolated by isopropanol precipitation, washed once with cold 100% Ethanol, air-dried and resuspended in TE low EDTA buffer. Any DNA contamination was eliminated by Turbo DNAse (Invitrogen #AM2238) reaction for 1h at 37°C. 500ng of RNA has been diluted in 250 μl of TE low EDTA buffer and applied to pre-equilibrated positively charged nylon membrane (Amersham Hybond-N+ # RPN203B) on Slot blot apparatus (Biorad # 1706542) by gentle vacuum. After one wash with 250 μl TE buffer, membrane was UV-crosslinked and blocked with 2% non-fat milk in PBS for 1h at room temperature. Then, primary m5C antibody (1:500) resuspended in PBS containing 2% milk and incubated overnight at 4°C. After secondary antibody (1:1000) incubation in 2% milk (PBS) for 1h at room temperature, RNA was visualized by autoradiography.

### Immunofluorescence

U2OS cells were reverse transfected with siNC and siNSUN2 (60nM). After 24h cells were transfected with FLAG-NSUN2 and FLAG-NSUN2-K190M. Then cells were seeded on poly-L-lysine coated coverslips to reach 60% confluency at the time of ionising irradiation. Cells were damage with 5 Gy and fixed (4% PFA in PBS for 10 min at 37°C) at certain time points. After permeabilization (0.2% Triton X-100 in PBS for 10 min at room temperature) and blocking (5% BSA in PBS for 1h at 37°C) steps, FLAG and Phospho-Histone H2Ax (Ser139) primary antibodies (1:1000) were incubated in 5% BSA solution overnight at 4°C. Then, coverslips were incubated with Alexa Fluor 488 (Invitrogen #A-11001) and Alexa Fluor 555 (Invitrogen #A-21428) secondary antibodies (1:2000) for 2h in dark, sealed with transparent nail varnish and visualized with Olympus FV1200 confocal microscope with 60X Oil Immersion Objective. Images were quantified using Cell profiler.

### Laser micro-irradiation

HEK 293T and HEK 293T 2B2 were treated with Doxycicline (1 μg/μl) for 3 days and transiently transfected with NeonGreen-NSUN2 or NeonGreen-NSUN2-K190M. Then, cells were seeded onto 35 mm glass bottom dishes (Greiner #627860) to reach the confluency of 60 % at the time of imaging. Before laser-induced damage, cells were sensitized by incubation (30 min at 37°C) with 10 μM Hoescht 33342 (Thermo Scientific #H3570). During laser micro-irradiation experiment cells were maintained in an incubation chamber at 37 °C and 5% CO_2_. Laser stripes were generated on a Nikon Spinning Disk SoRa microscope by a 405 pulsed laser in loops of 80 iterations and average peak powers of 1.5 mW and 2.4 mW. Recruitment and retention of NSUN2 at the site of induced damage was then monitored by tracking the fluorescence intensity at 488nm (1 frame every 4 seconds). Time-elapse images were analyzed using Fiji software.

### ChrRNA-Seq Data Processing

ChrRNA-Seq adapters were trimmed using Cutadapt (version 4.4) (https://cutadapt.readthedocs.io/en/stable/installation.html) in paired end mode and the quality of the resulting fastq files were assessed using FastQC (https://www.bioinformatics.babraham.ac.uk/projects/fastqc/). The trimmed reads were then aligned to human hg19 reference genome using STAR aligner ^44^. Each alignment file was then split using Samtools into two alignment files containing positively stranded and negatively stranded reads (https://www.htslib.org/)

### Metagene Plots

Strand specific coverage files containing CPM normalized read count per nucleotide position was generated for each alignment using deepTools bamCoverage (https://deeptools.readthedocs.io/en/develop/) ^45^. ComputeMatrix operation of deepTools was then performed on the strand separated bigwig files to calculate the CPM coverage in the 2.5kb flanking region of AsiSI site with bin size set at one. Bedtools intersect was used to find region of genes in 2.5kb flank of each annotated AsiSI site. Then custom python script was employed to annotate the bin values in the positively stranded matrix as sense or antisense based on whether they lie in same or opposite orientation of gene regions near AsiSI respectively. Similarly, the negatively stranded matrix was annotated as sense or antisense using the above logic. Only bins laying within gene regions were utilized for sense/antisense annotation. The sense matrices from positively and negatively stranded matrices were concatenated to form a combined sense matrix containing read coverage in sense orientation around AsiSI site. Antisense matrix was also crafted in the same manner to represent antisense matrix in 2.5kb flank region near each annotated AsiSI site. Antisense reads corresponding to different gene regions lying within same AsiSI site were summed to ensure that the matrix contains each AsiSI site as row with 5000 bins as columns representing antisense coverage in 2.5kb flank region of annotated AsiSI. Similar procedure was also followed to generate sense matrix. Matrix was then subdivided into different categories based on the known annotation of AsiSI sites as HR prone, NHEJ prone, uncut, highly transcriptionally active or transcriptionally less active sites. Sense and antisense matrix were then averaged across the AsiSI sites and plotted as line plots with separate scales using matplotlib python package. Fill plots representing read coverage in 2.5kb flank region of individual AsiSI sites were also created using matplotlib as replacement for IGV snapshots.

### PCA Plots

Bigwigsummary function of deeptools was employed in conjunction with plotPCA function to compare read coverage in 5kb region of BLESS 80 AsiSI sites between sample replicates.

### Box Plots

Coverage from sense and antisense matrices were used to make box plots representing CPM normalized reads in 500bp flank region of each AsiSI site. Coverage was calculated by summing CPM values in 500 bins centering DSB for each AsiSI site from sense and antisense matrix respectively. Box plots were then made using matplotlib python package and significance determined with 2 sample Wilcoxon Test from scipy python package. The comparison of CPM coverage between HR and NHEJ sites in 500bp flank region of AsiSI were performed using Mann Whitney U Test from scipy python package.

Fold changes across 500bp region flanking AsiSI were calculated by taking ratio of coverage in these regions between condition and control matrix for sense and antisense separately. Log2Fold Changes were then represented as box plots and significant difference in median values between sense and antisense were determined using 2-sample Wilcoxon test.

### Heatmaps

plotHeatmap function of deepTools was used to make the heatmaps with regions set as all annotated DSBs arranged in ascending order of cleavage efficiency. Separate heatmaps were generated using the above procedure for sense and antisense matrices.

### BLESS Seq data processing

BLESS Seq (E-MTAB-5817) was processed using the same protocol as detailed in ^30^. Read count coverage was calculated for all annotated DSBs (+-500bp) using bedtools multicov. The sites were then ordered based on read count coverage for representing cleavage efficiency of DSB sites.

### HR and NHEJ reporter assay

HeLa cells stably expressing EJ5-GFP and DR-GFP cassette were reversed transfected with 60nM of siNC, siNSUN2, siBRCA1. After 24h cells were seeded and forward transfected with pCBASceI plasmid (Addgene #26477)^46^ and Lipofectamine 3000. After 48h, cells were trypsinized and resuspended with 10%FBS in PBS before running FACS. siBRCA1 was used as positive control for HeLa HR reporter cells; the DNA-PK inhibitor Wortmannin (Sigma, W3144-250UL, 1µM for 6h) was used as the positive control for HeLa NHEJ reporter cells. Samples were acquired by CytoFLEX Flow cytometer (Beckman Coulter) and analyzed using FlowJo software.

### Comet assay

5000 Hela cells previously transfected with 60nM of siNC, siNSUN2 and siRAD51 were embedded in 0.5% CometAssay LMAgarose (Bio-techne #4250-050-02) before being spotted on Cometslides (Bio-techne #4250-050-03). On-gel cell lysis was performed by placing the Cometslide into lysis buffer (2.5M NaCl, 0.1M EDTA, 10mM Tris-Base, 10% DMSO, 1% TritonX-100, pH=10) overnight at 4°C. After wash with ddH_2_O, Cometslides were immersed into running buffer (0.3M NaOH, 1mM EDTA, pH=13) for 1h at 4°C followed by gel electrophoresis at 300mA for 30 min. Then, Cometslides were washed twice with Neutralization Buffer [0.4M Tris-base buffer (pH=7.5)] for 5min at room temperature, incubated with 70% ethanol for 15min and air dried. For chromatin tail visualization, incubation of 2μg/mL DAPI (BD Biosciences #564907) in PBS for 5min followed by 5min ddH_2_O washing was performed. Images were acquired by EVOS M7000 microscope with 10X objectives. To perform the tail moment quantification, Fiji with OpenComet plugin was used. Statistical significance was determined by using unpaired Welch’s correction.

### Western blot

Approximatively 1×10^7^ cells were lysed in 200 μl of RIPA buffer (Thermo Scientific #89901) supplemented with 1x Protease inhibitor (Roche # 11873580001) and incubated at 4°C for 20 min. Soluble supernatant was collected after centrifugation of 5 min at 500g (4°C), protein concentration was determined by Bradford colorimetric method (BioRad #5000006). Total proteins (20 μg) of whole cell extracts were resuspended in 1x LAemmli Buffer (BioRad #1610747), boiled for 10 min at 95°C and examined by western blot using 4-15% Mini-PROTEAN TGX precast protein gels (BioRad #4561083). After blocking step with 10% non-fat milk for 1h at room temperature, primary antibodies (1:1000) were incubated in 10% non-fat milk overnight at 4°C. Secondary antibodies (1:20000) were incubated for 1h at room temperature and blots were imaged with ECL substrate (Thermo Scientific #32106) and Amershan Hyperfilm ECL (Cytiva # 28-9068-35).

### MTS proliferation assay

For cellular viability analysis, 96-well plates were seeded with 5,000 HeLa cells previously transfected with siNC, siRAD51 and NSUN2 and damaged with 5 Gy. The MTS assay was carried out at certain time points using The CellTiter 96 AQueous One Solution Cell Proliferation Assay (Promega #G3580) according to the manufacturer’s instructions, and absorbance was measured at 490 nm.

### Statistical Analysis

Statistical tests were performed in GraphPad Prism. All error bars represent mean ± SD unless stated differently. The Kolmogorov-Smirnov normality test was performed to test for a normal distribution. If data meets normal distribution, statistical testing was performed using the Student’s *t*-test, one-way ANOVA, or unpaired Welch’s correction (for comet assay analysis). If data did not show a normal distribution, Mann–Whitney test for two groups (non-parametric comparison for PLA foci analysis), or Dunn’s test with Bonferroni corrections for multiple group comparisons. Significances are listed as \**p* ≤ 0.05, \*\**p* ≤ 0.01, \*\*\**p* ≤ 0.001, \*\*\*\**p* ≤ 0.0001.

## Data availability

Data reported in this paper can be shared by the lead contact upon request.

ChrRNA-seq data have been deposited to GEO and can be accessed under GSE260748 with private token: arubcusqrhabhit.

Any additional information required to re-analyze the data reported in this work paper is available from the lead contact upon request.

## Acknowledgements

We thank all members of Gullerova lab for their support. We are grateful to Dr Esra Balikci and Prof Kilian Huber for their help with preliminary experiments.

This work was supported by the Senior Research Fellowship by Cancer Research UK [grant number BVR01170], EPA Trust Fund [BVR01670], and Lee Placito Fund awarded to M.G.

## Author contributions

A.A. designed and performed most of the experiments and prepared initial draft of the manuscript. A.D.F. purified NSUN2 and DICER proteins. K.A. performed the bioinformatic analysis of Chr-RNA-seq. Q.L. performed comet, MTT and reporter assays. M.G. designed and supervised the project and wrote the manuscript. All the authors reviewed and approved the final version of the manuscript.

---

**Extended Data Fig. 1.**
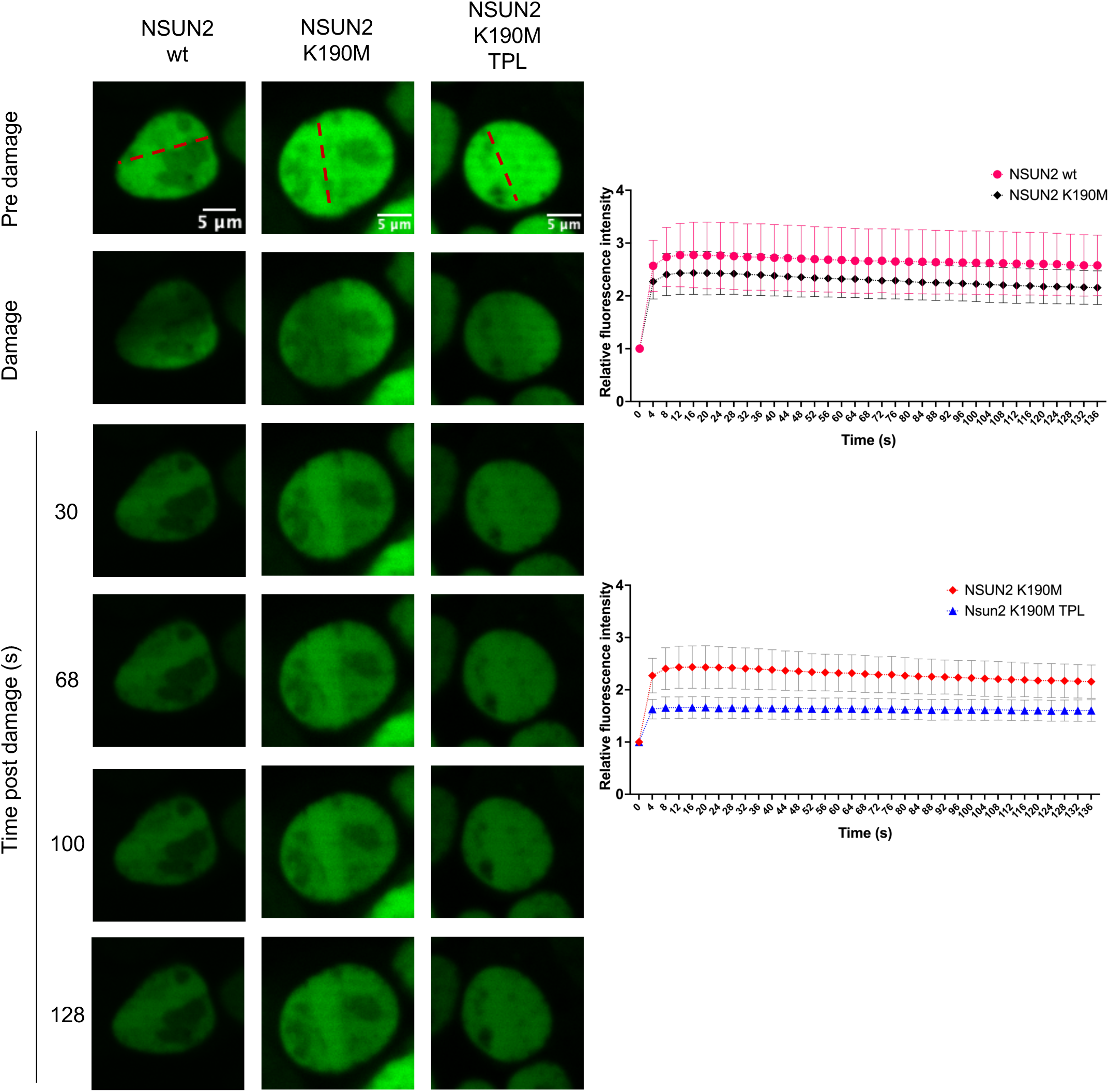
Laser induced DNA damage of HEK 293T cells transiently transfected with Neon-Green-NSUN2, Neon-Green-NSUN2 K190M plasmids and incubation with or without Triptolide (TPL, 1h 10μM). Left: Representative confocal microscopy images before and after the laser induced damage. Right: quantification of relative Neon-Green signal (background signal/ROI signal) (n>15) along the time considered (0-132 s). Error bars: mean ± SEM. Statistical significance was determined by two-way ANOVA with multiple comparison test (p ≤ 0.0001).

**Extended Data Fig. 2.**
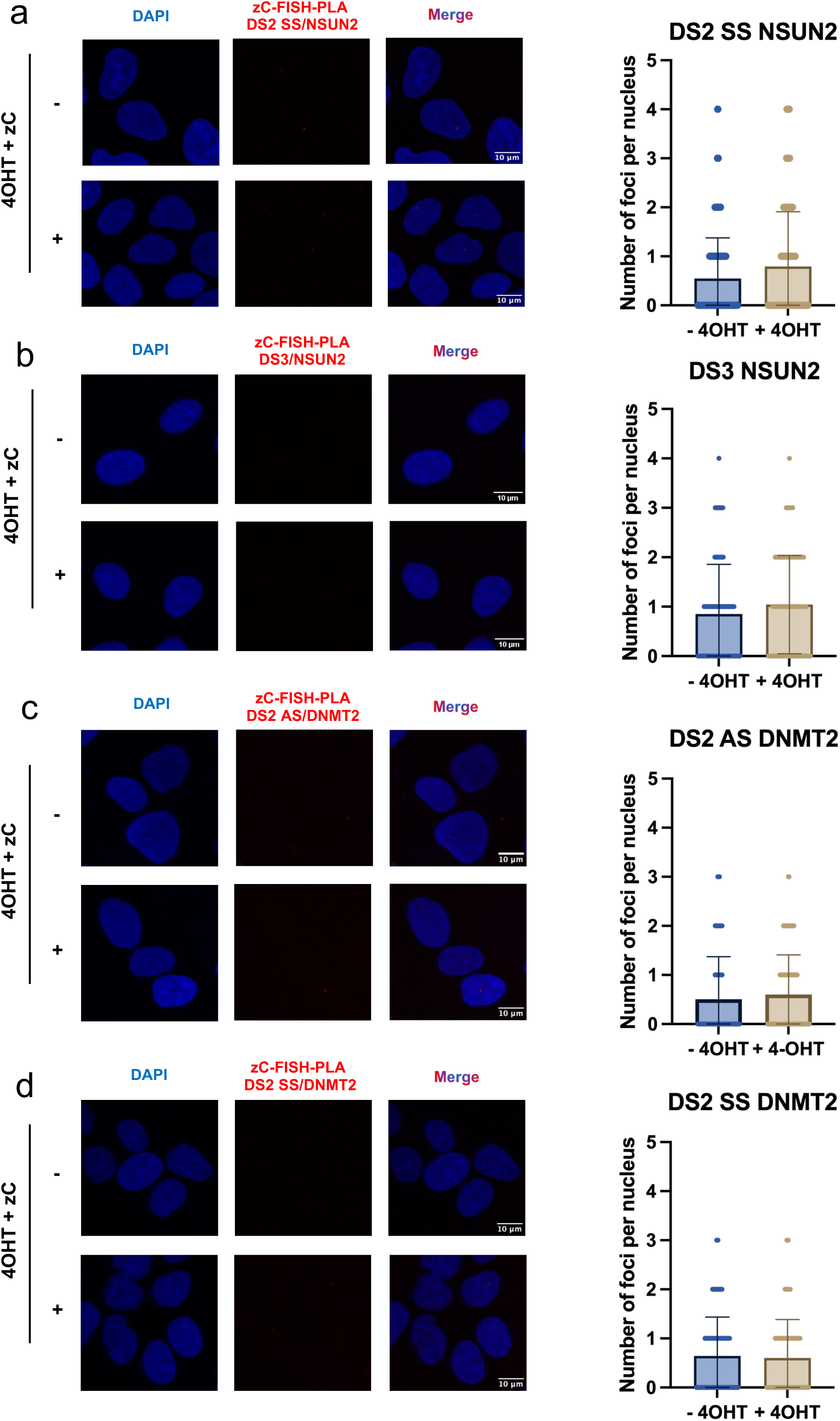
**a)** zC-FISH-PLA of NSUN2 and DNA probes (DS2 SS) with (+4OHT) or without (-4OHT) 4hr of (Z)-4-Hydroxytamoxifen. Quantification plot of FISH-PLA nuclear foci. Errors bars, mean ± SD. N>100. Statistical significance was determined using non-parametric Mann-Whitney test. Non-significant p-values not shown. **b)** zC-FISH-PLA of NSUN2 and DNA probes (DS3) with (+4OHT) or without (-4OHT) 4hr of (Z)-4-Hydroxytamoxifen. Quantification plot of FISH-PLA nuclear foci. Errors bars, mean ± SD. N>100. Statistical significance was determined using non-parametric Mann-Whitney test. Non-significant p-values not shown. **c)** zC-FISH-PLA of DNMT2 and DNA probes (DS2 AS) with (+4OHT) or without (-4OHT) 4hr of (Z)-4-Hydroxytamoxifen. Quantification plot of FISH-PLA nuclear foci. Errors bars, mean ± SD. N>100. Statistical significance was determined using non-parametric Mann-Whitney test. Non-significant p-values not shown. **d)** zC-FISH-PLA of DNMT2 and DNA probes (DS2 SS) with (+4OHT) or without (-4OHT) 4hr of (Z)-4-Hydroxytamoxifen. Quantification plot of FISH-PLA nuclear foci. Errors bars, mean ± SD. N>100. Statistical significance was determined using non-parametric Mann-Whitney test. Non-significant p-values not shown.

**Extended Data Fig. 3.**
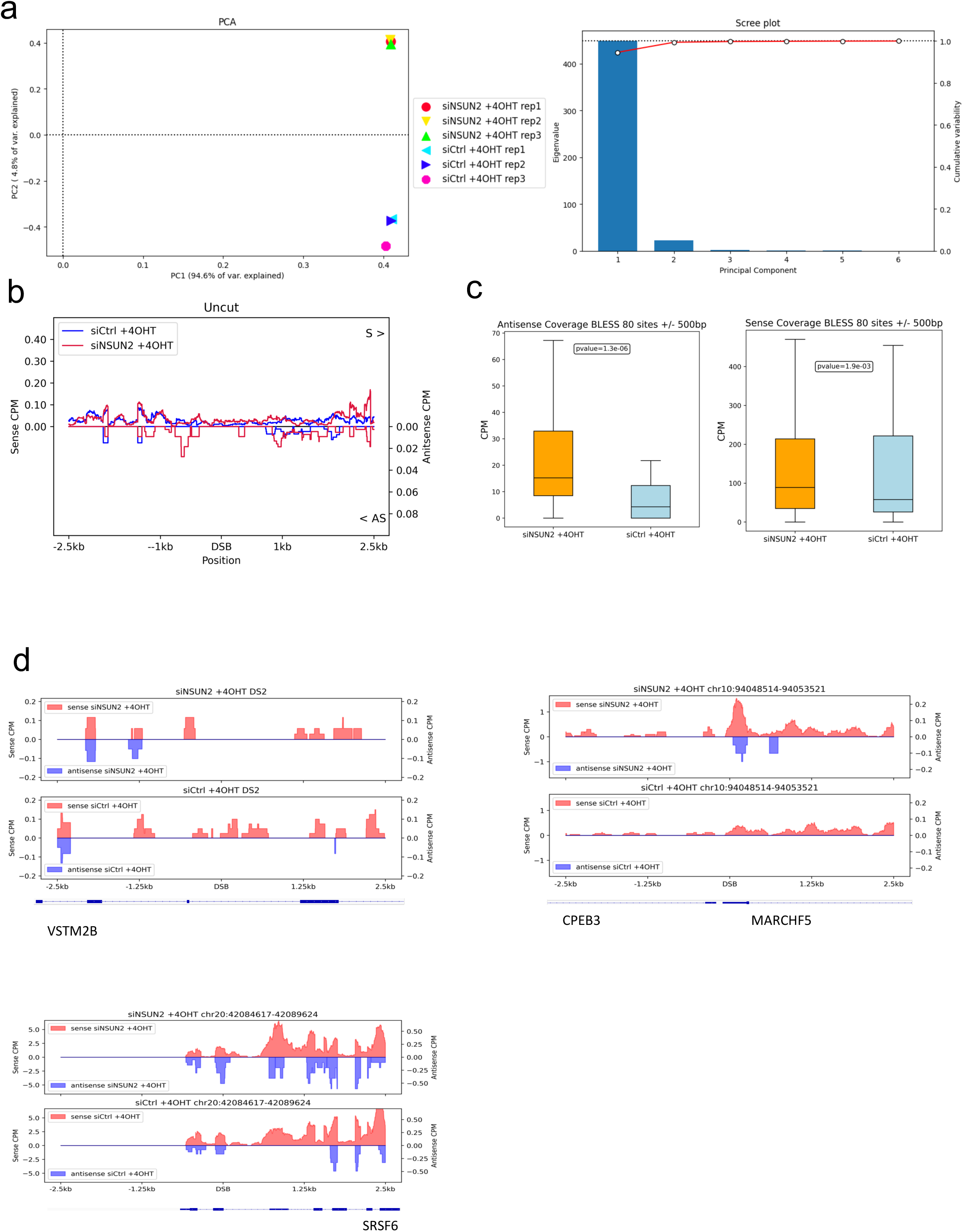
**a)** PCA plots show a comparison of chrRNA-Seq coverage in the 5kb flanking region of BLESS 80 *Asi*SI sites between sample replicates in NSUN2 knockdown and control conditions. **b)** Metagene plot shows chrRNA-Seq sense and antisense coverage in control (siCtrl) and INTS6 knockdown (siINTS6) cells with damage induction (+4-OHT) around 2.5kb flank region of uncut AsiSI sites (n=20). The reference genome is human hg19. **c)** Box plots show log2FoldChange of chrRNA-Seq coverage of sense reads and antisense reads upon NSUN2 knockdown in damage conditions compared to control for all AsiSI (+/- 500bp). Wilcoxon 2 sample test is used for statistical testing of medians between sense and antisense log2FoldChange distribution. **d)** Representative snapshots of individual genes showing sense and antisense chrRNA-Seq coverage in NSUN2 knockdown and control with damage induction around 2.5kb flank region of AsiSI cut. The specific loci information is listed on top of the snapshots. The reference genome is human hg19.

**Extended Data Fig. 4.**
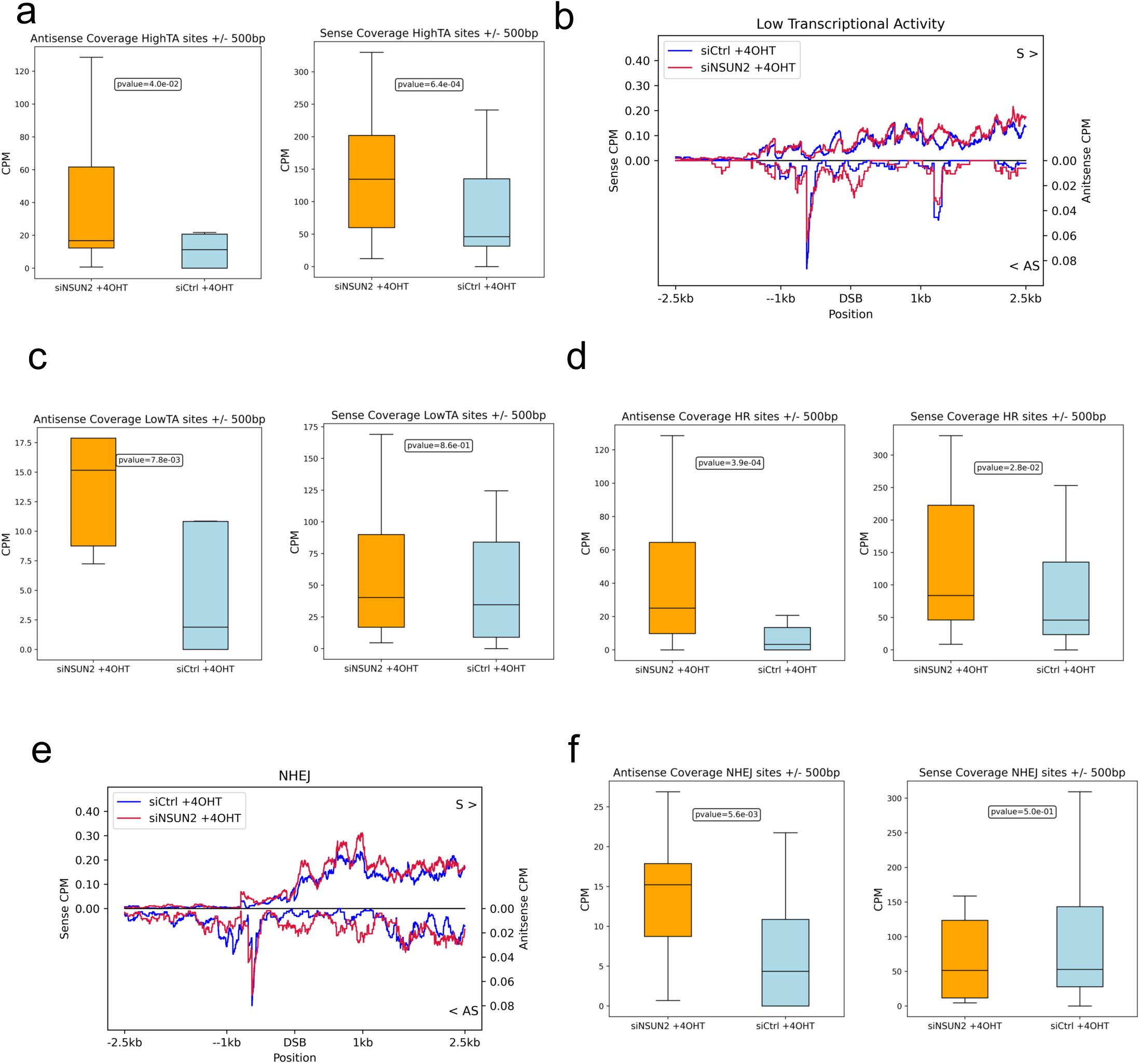
**a)** Box plots show log2FoldChange of chrRNA-Seq coverage of sense reads and antisense reads upon NSUN2 knockdown in damage conditions compared to control for AsiSI associated with highly transcribed regions (+/- 500bp). Wilcoxon 2 sample test is used for statistical testing of medians between sense and antisense log2FoldChange distribution. **b)** Metagene plot shows chrRNA-Seq sense and antisense coverage in control (siCtrl) and NSUN2 knockdown cells with damage induction (+4-OHT) around 2.5kb flank region of AsiSI sites associated with low transcription (n=80). The reference genome is human hg19. **c)** Box plots show log2FoldChange of chrRNA-Seq coverage of sense reads and antisense reads upon NSUN2 knockdown in damage conditions compared to control for AsiSI associated with low transcription (+/- 500bp). Wilcoxon 2 sample test is used for statistical testing of medians between sense and antisense log2FoldChange distribution. **d)** Box plots show log2FoldChange of chrRNA-Seq coverage of sense reads and antisense reads upon NSUN2 knockdown in damage conditions compared to control for HR prone AsiSI (+/- 500bp). Wilcoxon 2 sample test is used for statistical testing of medians between sense and antisense log2FoldChange distribution. **e)** Metagene plot shows chrRNA-Seq sense and antisense coverage in control (siCtrl) and NSUN2 knockdown cells with damage induction (+4-OHT) around 2.5kb flank region of NHEJ prone AsiSI sites (n=80). The reference genome is human hg19. **f)** Box plots show log2FoldChange of chrRNA-Seq coverage of sense reads and antisense reads upon NSUN2 knockdown in damage conditions compared to control for NHEJ prone AsiSI (+/- 500bp). Wilcoxon 2 sample test is used for statistical testing of medians between sense and antisense log2FoldChange distribution.

**Extended Data Fig. 5.**
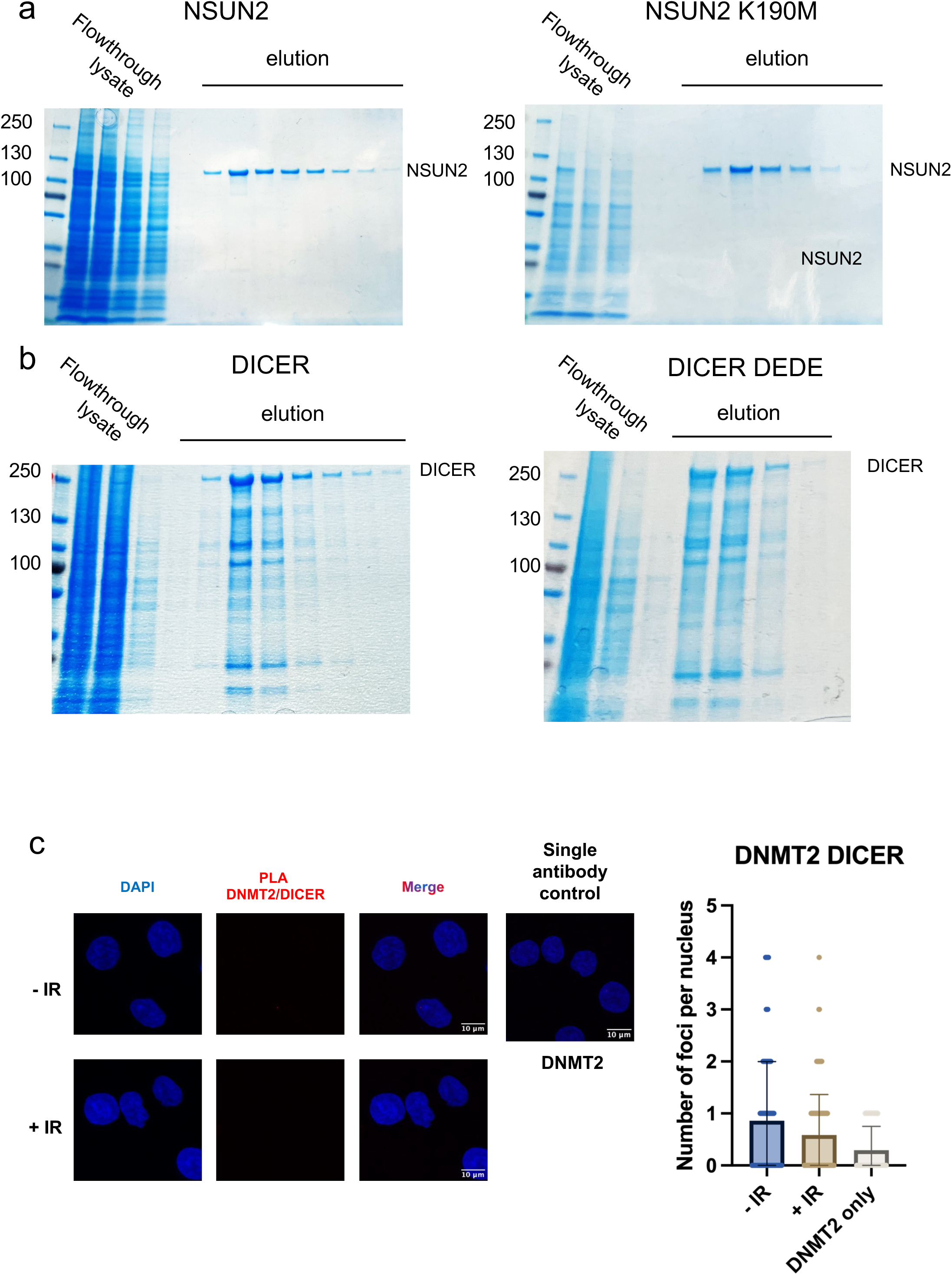
**a)** Coomassie Brilliant Blue staining of NSUN2 and NSUN2 K190M protein purification. **b)** Coomassie Brilliant Blue staining of DICER and DICER DEDE (D1320A; E1444A; D1709A; E1813A) protein purification. **c)** PLA of DNMT2 and DICER with (+IR) or without (-IR) Ionizing Radiation (10Gy, 15 min). As negative control single antibody assay was performed (DNMT2 only). Left: representative microscope images. Right panel: quantification of PLA nuclear foci. Errors bars, mean ± SD. N>100. Statistical significance was determined using non-parametric Mann-Whitney test. Non-significant p-values not shown.

**Extended Data Fig. 6.**
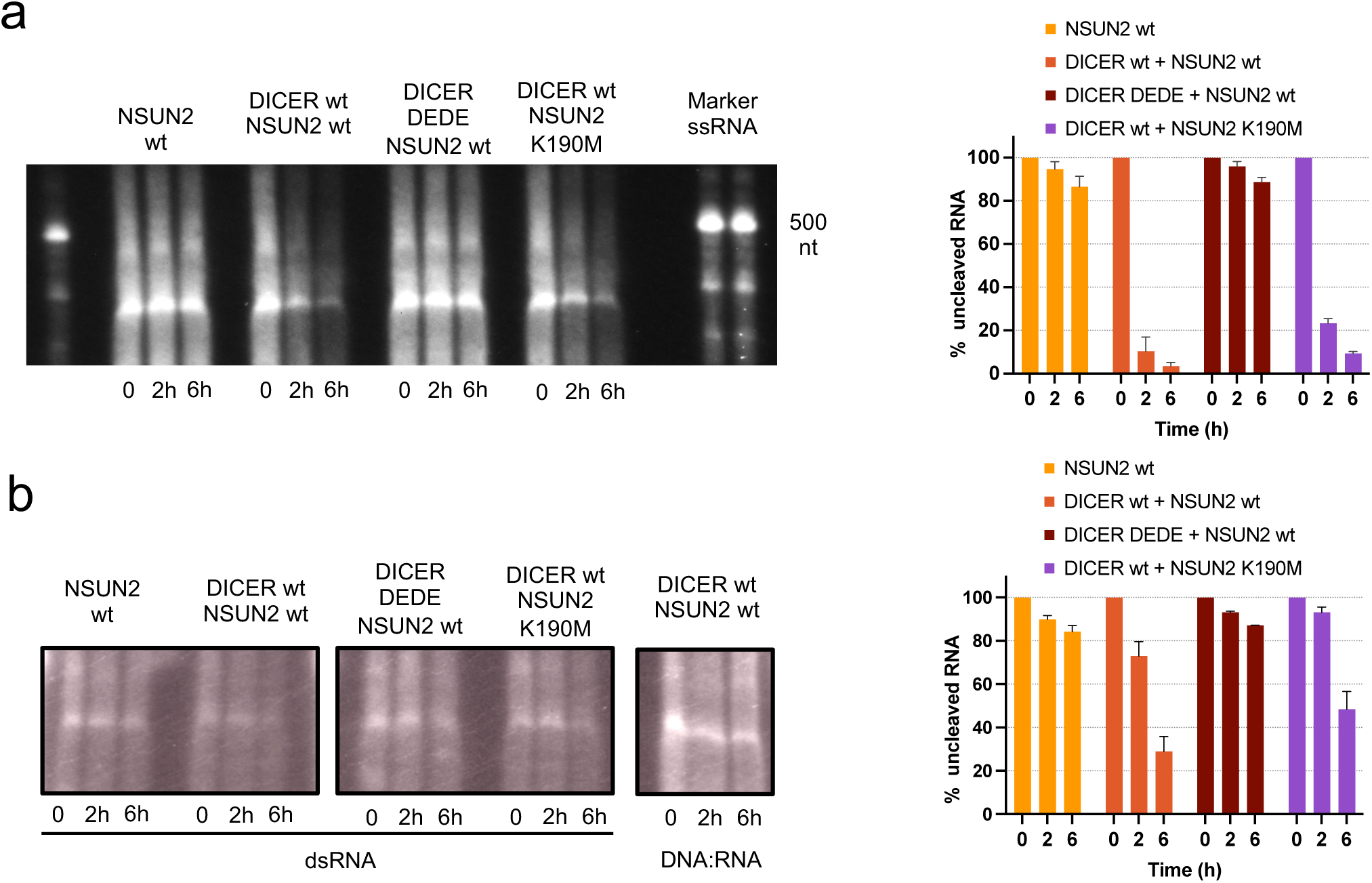
**a)** Northern blot of DICER cleavage reaction and in vitro transcribed (IVT) ssRNA DS2 derived after incubation with NSUN2 wt only, DICER wt and NSUN2, DICER DEDE and NSUN2 wt, DICER wt and NSUN2 K190M at certain time points (0, 2h, 6h). ssRNA Marker (Low Range ssRNA Ladder, NEB # N0364S). Quantification plot of cleaved ssRNA along the time points considered. Errors bars, mean ± SD. N=3. **b)** Northern blot of DICER cleavage reaction and IVT dsRNA and DNA:RNA heteroduplex DS2 derived after incubation with NSUN2 wt only, DICER wt and NSUN2, DICER DEDE and NSUN2 wt, DICER wt and NSUN2 K190M at certain time points (0, 2h, 6h). Quantification plot of cleaved dsRNA along the time points considered. Errors bars, mean ± SD. N=3.

**Extended Data Fig. 7.**
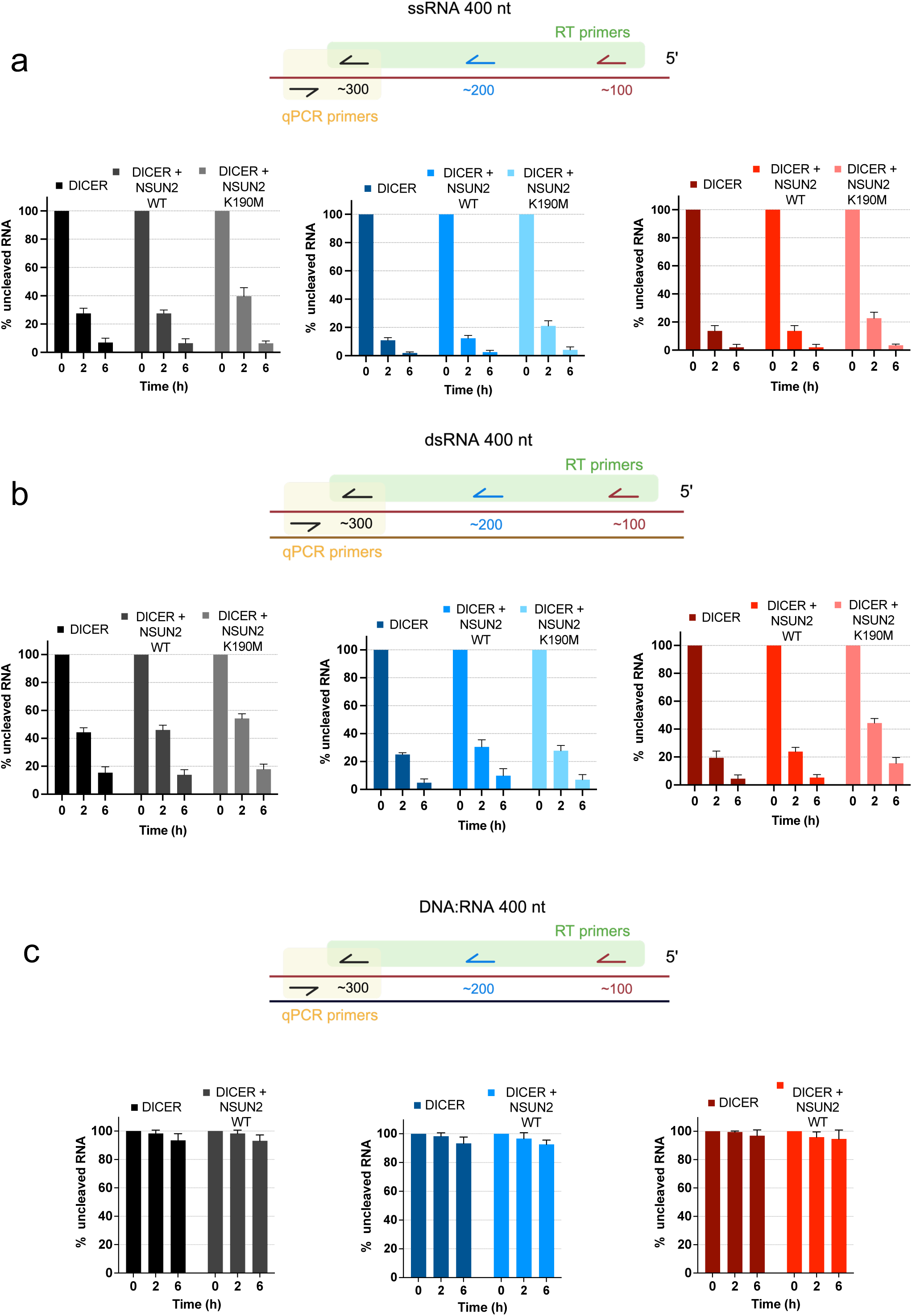
**a)** Top: Schematic representation of in vitro transcribed (IVT) antisense single stranded RNA DS2 derived with RT primers: (red arrow ∼100 nt, blue arrow ∼200 nt and black arrow ∼300nt away from the *Asi*Si site, highlighted in green), and qPCR primers (black arrows, highlighted in yellow). Bottom: quantification plots of cleaved RNA at 0, 2h and 6h time points in presence of DICER only, DICER + NSUN2 wt and DICER + NSUN2 K190M. Black, blue and red bars correspond to black, blue and red primer specific RT, respectively. Errors bars, mean ± SD. N=2. Statistical significance was determined using non-parametric Mann-Whitney test. Non-significant p-values not shown. **b)** Top: Schematic representation of IVT double stranded RNA DS2 derived with RT primers: (red arrow ∼100 nt, blue arrow ∼200 nt and black arrow ∼300nt away from the *Asi*Si site, highlighted in green), and qPCR primers (black arrows, highlighted in yellow). Bottom: quantification plots of cleaved RNA at 0, 2h and 6h time points in presence of DICER only, DICER + NSUN2 wt and DICER + NSUN2 K190M. Black, blue and red bars correspond to black, blue and red primer specific RT, respectively. Errors bars, mean ± SD. N=2. Statistical significance was determined using non-parametric Mann-Whitney test. Non-significant p-values not shown. **c)** Top: Schematic representation of DNA:RNA heteroduplex DS2 derived with RT primers: (red arrow ∼100 nt, blue arrow ∼200 nt and black arrow ∼300nt away from the *Asi*Si site, highlighted in green), and qPCR primers (black arrows, highlighted in yellow). Bottom: quantification plots of cleaved RNA at 0, 2h and 6h time points in presence of DICER only and DICER + NSUN2 wt. Black, blue and red bars correspond to black, blue and red primer specific RT, respectively. Errors bars, mean ± SD. N=2. Statistical significance was determined using non-parametric Mann-Whitney test. Non-significant p-values not shown.

**Extended Data Fig. 8.**
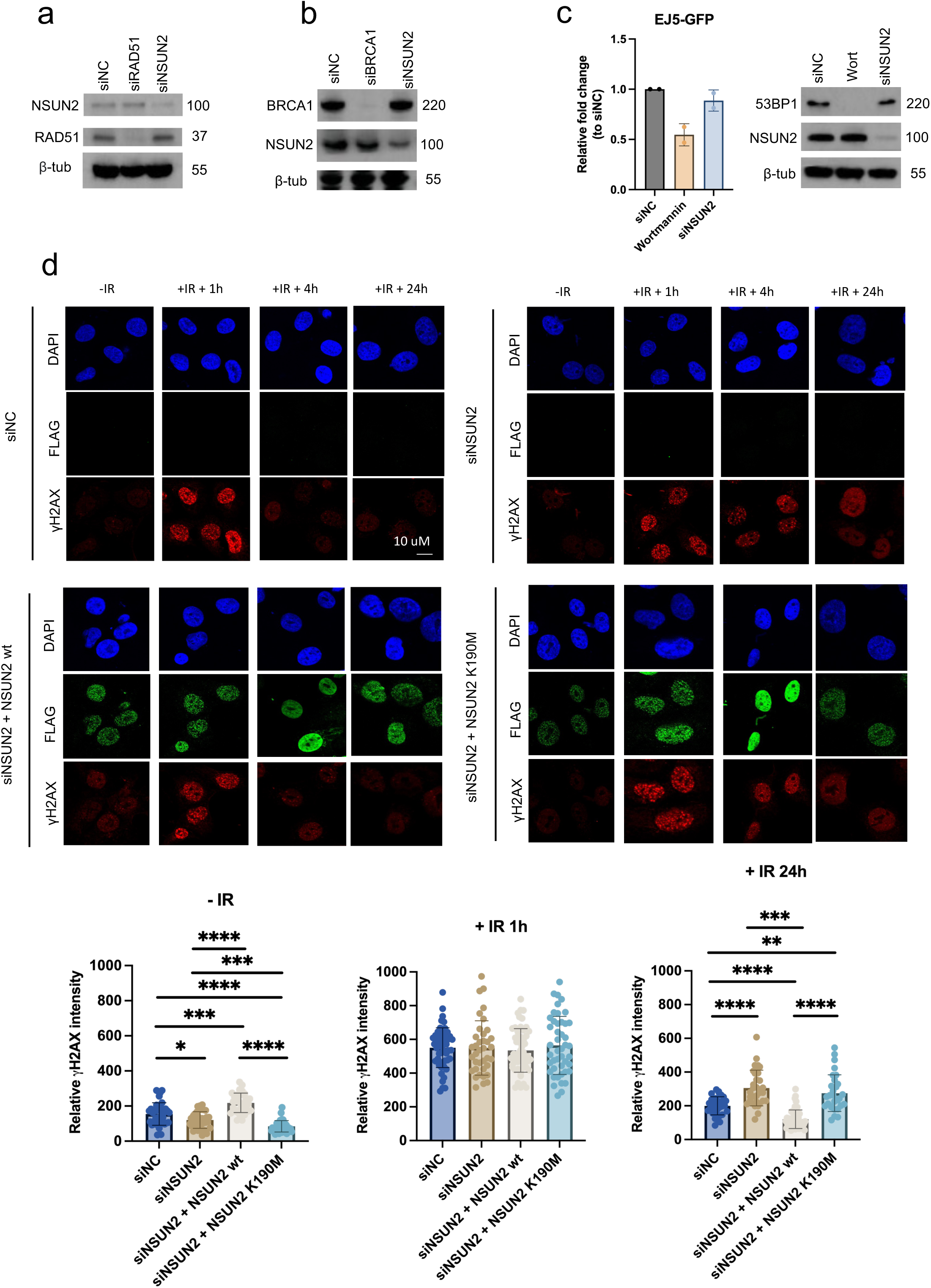
**a)** Western Blot showing the NSUN2 and RAD51 knockdown efficiency related to MTS assay. **b)** Western Blot showing the BRCA1 and NSUN2 knockdown efficiency related to DR-GFP reporter assay. **c)** Efficiency of NHEJ repair in EJ5-GFP reporter system in NSUN2 depleted (siNSUN2), siNegative Control (siNC) conditions and after incubation with Wortmannin. Errors bars, mean ± SD. Statistical significance was determined using non-parametric Mann-Whitney test. Non-significant p-values not shown. Right: Western Blot of the 53BP1 and NSUN2 knockdown efficiency related to EJ5-GFP reporter assay. **d)** Top: representative immunofluorescence images of FLAG-tagged NSUN2 wt, FLAG-tagged NSUN2 K190M and γH2Ax in NSUN2 depleted (siNSUN2), siNegative Control (siNC), NSUN2 wt rescued (siNSUN2 + NSUN2 wt) and NSUN2 K190M rescued (siNSUN2 + NSUN2 K190M) irradiated with (+IR +1h; +IR + 4h; +IR +24h) or without (-IR) 5 Gy. Bottom: Quantification plot of the γH2Ax intensity. Errors bars, mean ± SD. Statistical significance was determined using non-parametric Mann-Whitney test. \**p* ≤ 0.05, \*\**p* ≤ 0.01, \*\*\**p* ≤ 0.001, \*\*\*\**p* ≤ 0.0001.

